# Systematic analysis of dark and camouflaged genes: disease-relevant genes hiding in plain sight

**DOI:** 10.1101/514497

**Authors:** Mark T. W. Ebbert, Tanner D. Jensen, Karen Jansen-West, Jonathon P. Sens, Joseph S. Reddy, Perry G. Ridge, John S. K. Kauwe, Veronique Belzil, Luc Pregent, Minerva M. Carrasquillo, Dirk Keene, Eric Larson, Paul Crane, Yan W. Asmann, Nilufer Ertekin-Taner, Steven G. Younkin, Owen A. Ross, Rosa Rademakers, Leonard Petrucelli, John D. Fryer

**Author notes:** Contributed equally (Mark T. W. Ebbert, Tanner D. Jensen).

## Abstract

**Background:** The human genome contains ‘dark’ gene regions that cannot be adequately assembled or aligned using standard short-read sequencing technologies, preventing researchers from identifying mutations within these gene regions that may be relevant to human disease. Here, we identify regions that are ‘dark by depth’ (few mappable reads) and others that are ‘camouflaged’ (ambiguous alignment), and we assess how well long-read technologies resolve these regions. We further present an algorithm to resolve most camouflaged regions (including in short-read data) and apply it to the Alzheimer’s Disease Sequencing Project (ADSP; 13142 samples), as a proof of principle.

**Results:** Based on standard whole-genome Illumina sequencing data, we identified 37873 dark regions in 5857 gene bodies (3635 protein-coding) from pathways important to human health, development, and reproduction. Of the 5857 gene bodies, 494 (8.4%) were 100% dark (142 protein-coding) and 2046 (34.9%) were ≥5% dark (628 protein-coding). Exactly 2757 dark regions were in protein-coding exons (CDS) across 744 genes. Long-read sequencing technologies from 10x Genomics, PacBio, and Oxford Nanopore Technologies reduced dark CDS regions to approximately 45.1%, 33.3%, and 18.2% respectively. Applying our algorithm to the ADSP, we rescued 4622 exonic variants from 501 camouflaged genes, including a rare, ten-nucleotide frameshift deletion in *CR1*, a top Alzheimer’s disease gene, found in only five ADSP cases and zero controls.

**Conclusions:** While we could not formally assess the *CR1* frameshift mutation in Alzheimer’s disease (insufficient sample-size), we believe it merits investigating in a larger cohort. There remain thousands of potentially important genomic regions overlooked by short-read sequencing that are largely resolved by long-read technologies.

## Background

Researchers have known for years that large, complex genomes, including the human genome, contain ‘dark’ regions—regions where standard high-throughput short-read sequencing technologies cannot be adequately assembled or aligned—thus preventing our ability to identify mutations within these regions that may be relevant to human health and disease. Some dark regions are what we term ‘dark by depth’ (few or no mappable reads), while others are what we term ‘dark by mapping quality’ (reads aligned to the region, but with a low mapping quality). Regions that are dark by depth may arise because the region is inherently difficult to sequence at the chemistry level (e.g., high GC content [1, 2]), essentially eliminating sequencing reads from that region altogether. Other dark regions arise, not because the sequencing is inherently problematic, but because of bioinformatic challenges. Specifically, many dark regions arise from duplicated genomic regions, where confidently aligning short reads to a unique location is not possible; we term these regions as ‘camouflaged’. These camouflaged regions are generally either large contiguous tandem repeats (e.g., centromeres, telomeres, and other short tandem repeats), or a larger specific DNA region that has been duplicated (e.g., a gene duplication) either in tandem or in a more distal genome region. In fact, many genes in the human genome were duplicated over evolutionary time and are still transcriptionally and translationally active (e.g., heat-shock proteins) [3-9], while others have been duplicated, but are considered inactive (i.e., pseudogenes). Regardless of whether the duplication is active, however, any genomic region that has been nearly-identically duplicated, and is large enough to prevent sequencing reads from aligning unambiguously will be ‘dark’, because the aligner cannot determine which genomic region the read originated from.

When confronted with a read that aligns equally well to two or more camouflaged regions (commonly known as multi-mapping reads [2,10]), standard next-generation sequence aligners, such as the Burrows-Wheeler Aligner (BWA) [11-13], randomly assign the read to one of the regions and assign a low mapping quality. For BWA, specifically, reads that cannot be uniquely mapped are generally assigned a mapping quality (MAPQ) of 0; though, in certain paired-end sequencing scenarios, BWA will assign a high mapping quality if the read mate is confidently mapped nearby (i.e., within the estimated insert-size length).

Recent work has characterized camouflaged regions, in part, including a study that demonstrates how this issue affects all standard RNA-Seq analyses [10], and another that quantifies the number of nucleotides in human reference GRCh38 that are dark for mapping quality of 0 (camouflaged regions), based on 1000 Genome Project data [2]. Robert and Watson demonstrated that expression for 958 genes were either over- or under-represented because of multi-mapping reads across 12 different RNA-Seq processing methods, and no method was immune to the problem [10]. They also demonstrated that many of these genes are directly implicated in human disease. Zheng-Bradley et al. recently re-aligned genomes from the 1000 Genomes Project to GRCh38, and, among other findings, generally demonstrated the breadth of multi-mapping reads across the genome [2]. These data characterize the general problem, and report specific genes affected by this issue.

Here, we systematically analyze dark and camouflaged genes to more fully characterize the problem, and we highlight many disease-relevant genes that are directly implicated in Alzheimer’s disease, autism spectrum disorder, amyotrophic lateral sclerosis (ALS), spinal muscular atrophy (SMA), and others. We also show that long-read sequencing technologies substantially reduce the number of dark and camouflaged regions, and we present a method to address camouflaged regions, even in standard short-read sequencing data. As a proof of concept, we apply our method to the Alzheimer’s Disease Sequencing Project (ADSP) data, and identify a rare, ten-nucleotide frameshift deletion in the C3b and C4b binding domain of *CR1*, a top Alzheimer’s disease gene [14-22], that is only present in five ADSP cases and zero controls. The ADSP is not large enough to statistically assess association between the *CR1* frameshift mutation and Alzheimer’s disease.

## Results

To quantify the number of dark and camouflaged regions in standard short-read whole-genome sequencing data, we obtained whole-genome sequencing data for ten unrelated males from the Alzheimer’s Disease Sequencing Project (ADSP) and scanned each sample for dark and camouflaged regions, averaging across all ten samples; we only used data from males for this study so we could also assess dark and camouflaged regions on the Y chromosome because large portions of the Y chromosome are dark. We ignored incomplete genomic regions (e.g., centromeres). We then limited the dark and camouflaged regions to known gene bodies, based on annotations from build 87 of the Ensembl GRCh37 human reference genome [23]. All ten samples were sequenced using standard Illumina whole-genome sequencing with 100-nucleotide read lengths, where median genome-wide read depths ranged from 35.4x to 42.9x coverage, with an overall median of 39.4x. We performed the same analyses on ten unrelated males from the 1000 Genomes Project [24] that were sequenced using Illumina whole-genome sequencing with 250-nucleotide read lengths, where median genome-wide read depths ranged from 39.3x to 52.6x coverage, with an overall median of 48.9x. Similarly, we assessed how well long-read sequencing technologies, including 10x Genomics (52x median coverage), PacBio (50x median coverage), and ONT (46x median coverage) resolve dark and camouflaged regions. Although we were only able to obtain a single high-depth male genome for each long-read technology, we believe our results are a reasonable estimate for how well each technology addresses dark and camouflaged regions. Larger sequencing studies will further clarify our results.

We consider a region ‘dark’ for one of two reasons: (1) insufficient number of reads aligned to the genomic region (dark by depth); and (2) reads aligned to the region, but with insufficient mapping quality for a variant caller to identify mutations in the region (dark by mapping quality). Specifically, we define regions that are dark by depth as those with fewer than five aligned reads (Figure 1a), and regions that are dark by mapping quality as those where ≥90% of aligned reads have a mapping quality (MAPQ) <10 (Figure 1b). Defining dark-by-depth regions as those with fewer than five reads is a relatively strict cutoff, and likely underestimates the number of dark regions because 20 to 30 reads is often considered a reasonable minimum to confidently identify heterozygous mutations; overall median read depth is an important factor, however, and we believe a strict cutoff provides a more conservative estimate. We used a mapping quality threshold <10 to define regions that are dark by mapping quality because that is the standard cutoff used in the Genome Analysis ToolKit (GATK) [25]. Camouflaged regions are those that are dark by mapping quality because the region has been duplicated in the genome (Figure 1c). We identified sets of camouflaged regions (regions camouflaged by each other) using BLAT [26], where we required at least 98% sequence identity for two regions to be included in the same set.

**Figure 1.**
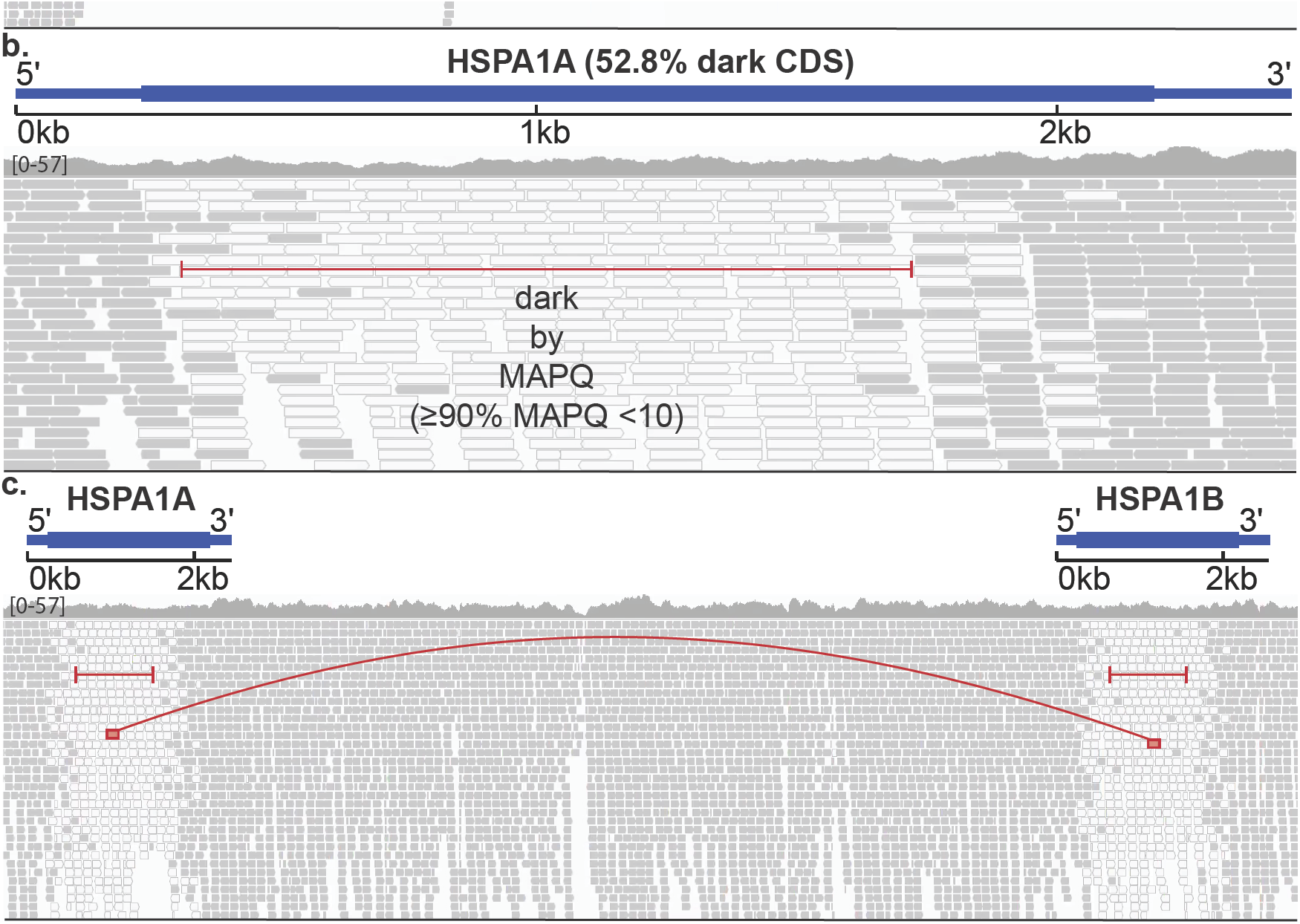
Genomic regions may be ‘dark’ by depth or mapping quality, many of which are ‘camouflaged’. Large, complex genomes are known to contain ‘dark’ regions where standard high-throughput short-read sequencing technologies cannot be adequately assembled or aligned. We split these dark regions into two types: (1) dark because of low depth; and (2) dark because of low mapping quality (MAPQ), which are mostly ‘camouflaged’, **(a)** *HLA-DRB5* encodes a Major Histocompatibility Complex protein that plays an important role in immune-response and has been associated with several diseases, including Alzheimer’s disease. It is well known to be dark (low depth); specifically, when performing whole-genome sequencing using standard short-read sequencing technologies, an insufficient number of reads align, preventing variant callers from assessing mutations. We calculated sequencing depth across *HLA-DRB5* for ten male samples from the Alzheimer’s Disease Sequencing Project (ADSP) that were sequenced using standard lllumina whole-genome sequencing with 100-nucleotide read lengths. Approximately 62.0% (50.2% of coding sequence) of *HLA-DRB5* is dark by depth (<5 aligned reads; indicated by red lines), **(b)** *HSPA1A* is a heat-shock protein from the 70-kilodalton (kDa) heat-shock protein family, and plays an important role in stabilizing proteins against aggregation. *HSPA1A* is dark because of low mapping quality (MAPQ <10 for ≥90% of reads at a given position). Approximately 41.8% (52.8% coding sequence) of *HSPA1A* is dark by mapping quality (indicated by red line). Dark gray bars indicate sequencing reads with a relatively high mapping quality, whereas white bars indicate reads with a low mapping quality (MAPQ = 0). **(c)** Many genomic regions that are dark because of mapping quality arise because they have been duplicated in the genome, which we term ‘camouflaged’ (or ‘camo genes’). When confronted with a read that aligns equally well to more than one location, standard sequence aligners randomly assign the read to one location and give it a low mapping quality. Thus, it is unclear from which gene any of the reads indicated by white bars originated from. *HSPA1A* and *HSPA1B* are clear examples of camouflaged genes arising from a tandem duplication. The two genes are approximately 14kb apart and approximately 50% of the genes are identical.

### Standard short-read sequencing leaves 37873 dark regions across 5857 gene bodies, including protein-coding exons from 744 genes

Using whole-genome Illumina sequencing data (100-nucleotide read lengths) from ten unrelated males, we identified 37873 dark regions (>16 million nucleotides) in 5857 gene bodies (based on Ensemble GRCh37 build 87 gene annotations) that were either dark by depth or dark by mapping quality (Supplemental Figure 1a; Supplemental Tables 1-2). Stratifying the gene bodies by GENCODE biotype [27], 3635 gene bodies were protein coding, 1102 were pseudogenes, and 720 were long intergenic non-coding RNAs (lincRNA; Figure 2a). Of all 37873 dark gene-body regions, 28598 were intronic, 4113 were in non-coding RNA exons (e.g., lincRNAs and pseudogenes), 2657 were in protein-coding exons (CDS), 1134 were in 3’UTR regions, and 1103 were in 5’UTR regions (Figure 2b; Supplemental Table 1). Any dark region that spanned a gene element boundary (e.g., intron to exon) was split into separate dark regions. Of the 5857 gene bodies, 494 (8.4%) were 100% dark, 1560 (26.6%) were at least 25% dark, and 2046 (34.9%) were at least 5% dark (Supplemental Figure 1b; Supplemental Table 1).

**Figure 2.**
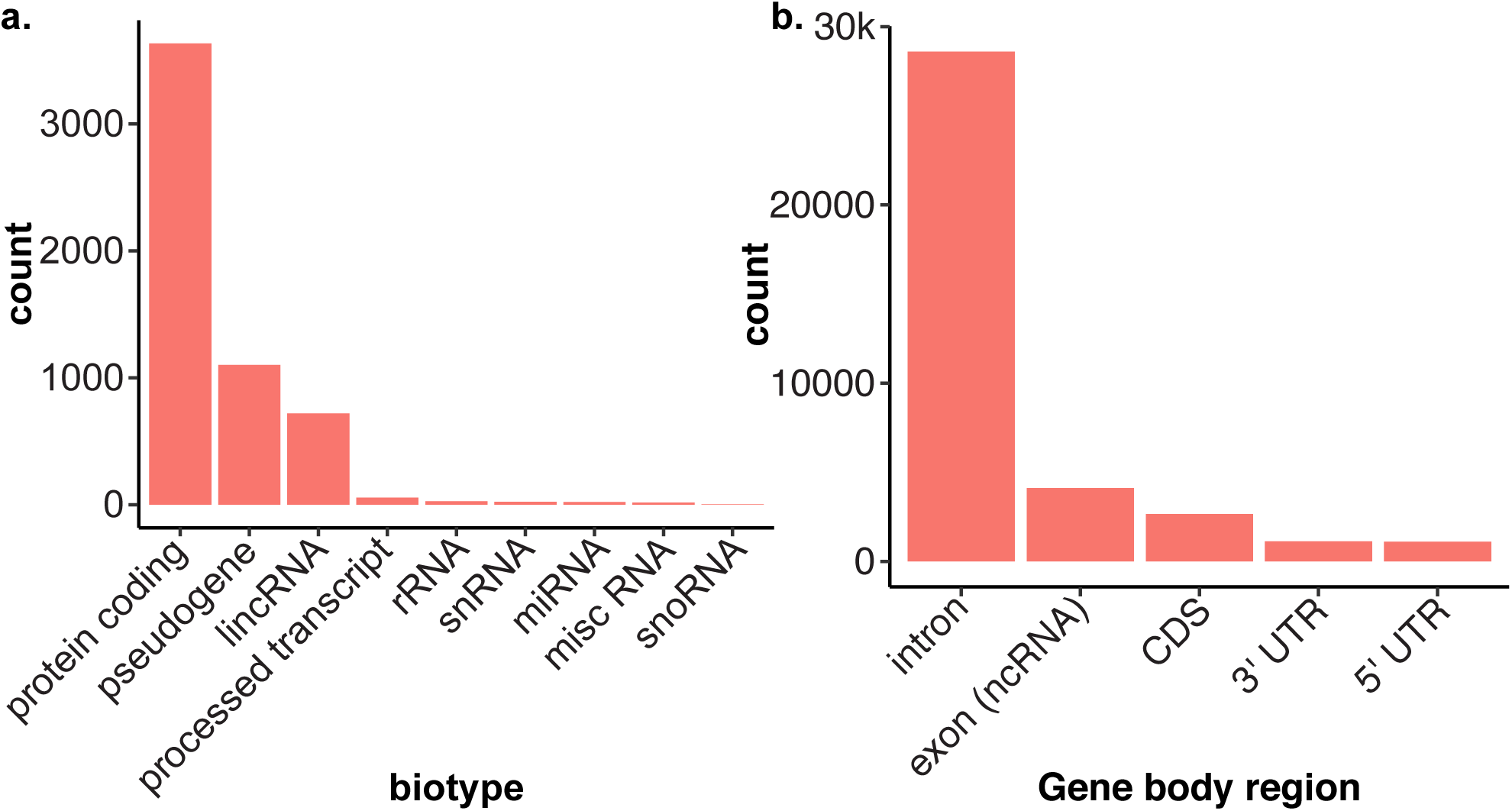
Many dark regions involve protein-coding gene regions. We identified 37873 dark regions (>16 million nucleotides) in 5857 gene bodies that were either dark by depth or dark by mapping quality, **(a)** Stratifying the gene bodies by GENCODE biotype, 3635 gene bodies were protein coding, 1102 were pseudogenes, and 720 were long intergenic non-coding RNAs (lincRNA). **(b)** Of all 37873 dark regions, 28598 were intronic, 4114 were in lincRNA exons, 2657 were in protein-coding exons (CDS), 1134 were in 3’UTR regions, and 1103 were in 5’UTR regions. Any dark region that spanned a gene element boundary (e.g., intron to exon) was split into separate dark regions.

Focusing only on CDS regions, we identified 2757 dark CDS regions (>460000 nucleotides) across 744 protein-coding genes that were dark by either depth or mapping quality (Figure 3a; Supplemental Tables 1-2). Exactly 142 (19.1%) of the 744 protein-coding genes were 100% dark in CDS regions, 441 (59.3%) were at least 25% dark in CDS regions, and 628 (84.4%) were at least 5% dark in CDS regions (Figure 3b; Supplemental Table 1). Exactly 474 of the 628 genes that were 5% dark in CDS regions were dark because they were camouflaged.

**Figure 3.**
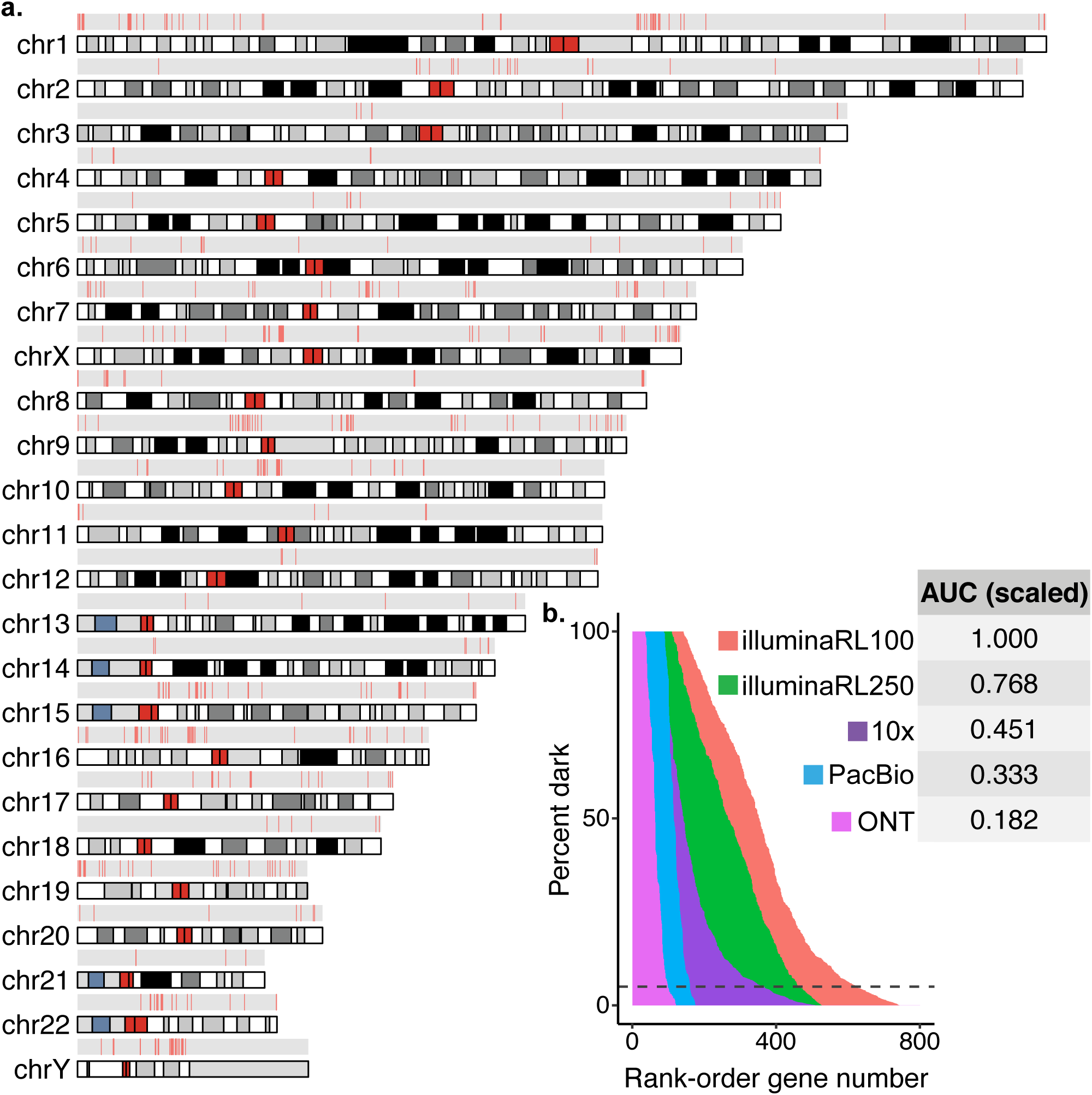
Dark coding regions occur throughout the genome, and are largely resolved with long-read sequencing technologies. We identified 2757 dark coding (CDS) regions (>460000 nucleotides) in 744 protein-coding genes that were dark by either depth or mapping quality (Supplemental Tables 1-2). Exactly 142 (19.1%) of the 744 protein-coding genes were 100% dark in CDS regions, 441 (59.3%) were at least 25% dark in CDS regions, and 628 (84.4%) were at least 5% dark in CDS regions (Supplemental Table 1). **(a)** We mapped all protein-coding gene bodies with a dark coding exon to the genome to visualize their genomic location, and are generally spread throughout. There are several tight clusters of dark CDS regions on chromosomes 1, 9,10, and Y, however. **(b)** We assessed how well increasing read lengths would resolve dark regions by assessing samples sequenced with lllumina whole-genome sequencing using 250-nucleotided read lengths, as well as long-read technologies 10x Genomics, Oxford Nanopore Technologies (ONT), and Pacific Biosciences (PacBio). Data from the samples sequenced using 250-nucleotide lllumina read lengths reduced the area under the curve by 23.2% in CDS regions; this translates to a 24.4% reduction in dark CDS nucleotides. Comparing long-read sequencing technologies to the standard lllumina 100-nucleotide read lengths, 10x Genomics, PacBio, and ONT reduced the area under the curve for CDS regions by approximately 54.9%, 66.7%, and 81.8%, respectively; this translates to a 57.8%, 68.6%, and 81.4% reduction in dark CDS nucleotides, respectively. The area under the curve (AUC) for each technology is scaled in reference to lllumina sequencing based on 100-nucleotide read lengths (i.e., AUC for lllumina 100-nucleotide read lengths = 1). In contrast to overall results, PacBio outperformed 10x Genomics when looking only at CDS regions (see text). Most analyses focused on genes where at least 5% of the CDS nucleotides are dark, indicated by the dashed line.

### Most dark regions are specifically camouflaged

Regions may be dark because of either low depth or low mapping quality, but the majority of regions are dark because of mapping quality, and specifically because they are camouflaged (low mapping quality because of a duplication). Exactly 3953 of the 5857 dark gene bodies are dark because of mapping quality, where 3252 are, in fact, camouflaged. We also measured the number of times each gene region was duplicated and found that 70% of gene regions were replicated three or fewer times in the genome, but 84 regions were duplicated ≥ 100 times (Supplemental Figure 2a), with the most repeated regions (ten separate intronic regions totaling 2235 nucleotides from *C5orf48*) being replicated 941 times in aggregate. Limiting to only CDS regions, we estimate that 74.1% are replicated three or fewer times, with 38 replicated ≥ 10 times (Supplemental Figure 2b) and the most repeated region was from *NBPF12*, in which 173 nucleotides were replicated 37 times.

### Long-read sequencing technologies resolve substantial portions of the dark regions

Data from the samples sequenced using 250-nucleotide Illumina read lengths reduced the percentage of dark nucleotides by 30.1% and 24.4% for all gene bodies, and for only CDS regions, respectively, leaving 69.9% and 75.6% of the nucleotides dark, respectively (Supplemental Figure 1b; Figure 3b; Supplemental Tables 3-4). Comparing long-read sequencing technologies to the standard Illumina 100-nucleotide read lengths, the ONT platform performed best, both when assessing entire gene bodies, and when considering only CDS regions. Specifically, approximately 41.2%, 25.8%, and 24.9% of the nucleotides remained dark for all gene bodies for PacBio, 10x Genomics, and ONT, respectively (Supplemental Figure 1b; Supplemental Tables 5-10). Similarly, approximately 42.2%, 31.4%, and 18.5% of CDS nucleotides remained dark for 10x Genomics, PacBio, and ONT, respectively (Figure 3b; Supplemental Tables 5-10). In contrast to overall gene-body results, PacBio outperformed 10x Genomics when looking only at CDS regions (Supplemental Figure 1b; Figure 3b). The long-read technologies improved over Illumina mostly by reducing the percentage of nucleotides that are dark by mapping quality (Supplemental Figure 1c). Surprisingly, the percentage of gene-body regions that are dark because of low depth is higher for all long-read technologies than it is for Illumina (Supplemental Figure 1c).

We generated a density plot for the length of all dark-by-mapping quality regions to approximate the proportion of regions each sequencing technology should be able to resolve (Supplemental Figure 3), which resulted in a bimodal distribution. The two modes are located at 95 and 538 nucleotides. As expected, median read lengths for the Illumina whole-genome sequencing based on 100-nucleotide and 250-nucleotide read lengths were 100 and 250 nucleotides, respectively. The first mode for the camouflaged region lengths is at 95, explaining why 100-nucleotide read lengths are insufficient to unambiguously span most dark-by-mapping quality regions. The 250-nucleotide read lengths fall between the two modes, explaining why 250-nucleotide read lengths resolve a high percentage of camouflaged regions. In other words, 100-nucleotide read lengths are too short to bridge most camouflaged regions, but 250-nucleotide read lengths appear to be sufficient for many. Median read lengths for both the ONT and PacBio genomes we used in this study were 6276 (N50 = 33973) and 8511 (N50 = 17467) nucleotides, respectively, which is shorter than expected, but substantially longer than necessary to resolve most camouflaged regions. We believe comparing median read lengths, rather than N50, is more useful in this scenario, because we are interested to know what percentage of reads are likely to bridge a given dark or camouflaged region. Our results suggest that our estimates for the percentage of camouflaged regions ONT and PacBio are able to resolve may be conservative because a longer DNA library should resolve even more camouflaged regions.

### Important pathways and gene families are affected by dark and camouflaged regions

Because such a large number of genes are dark, we characterized the pathways for genes that are not fully represented in standard Illumina short-read sequencing (100-nucleotide reads) datasets. We included all genes where at least 5% of the CDS regions were dark (670 unique gene symbols) and identified several pathways that are important in human health, development, and reproductive function (Figure 4a; Supplemental Table 11). Specific pathways included defensins (R-HSA-1461973; logP = −7.04), gonadal mesoderm development (G0:0007506; logP = −6.18), base-excision repair (G0:0006284; logP = −5.93), chromatin silencing (G0:0006342; logP = −5.86), Deubiquitination (R-HSA-5688426; logP = −5.32), NLS-bearing protein import into nucleus (G0:0006607; logP = −5.31), spindle assembly (G0:0051225; logP = −5.19), spermatogenesis (G0:0007283; logP = −4.93), and forebrain neuron differentiation (G0:0021879; logP = −4.09). Some specific gene families involved in these pathways include eleven defensin genes (e.g., DEFA1 and DEFB4A), five testis specific proteins (e.g., TSPY2), eleven ubiquitin-specific 17-like family members, and twelve golgin genes (e.g., GOLGA6B; Supplemental Table 11).

**Figure 4.**
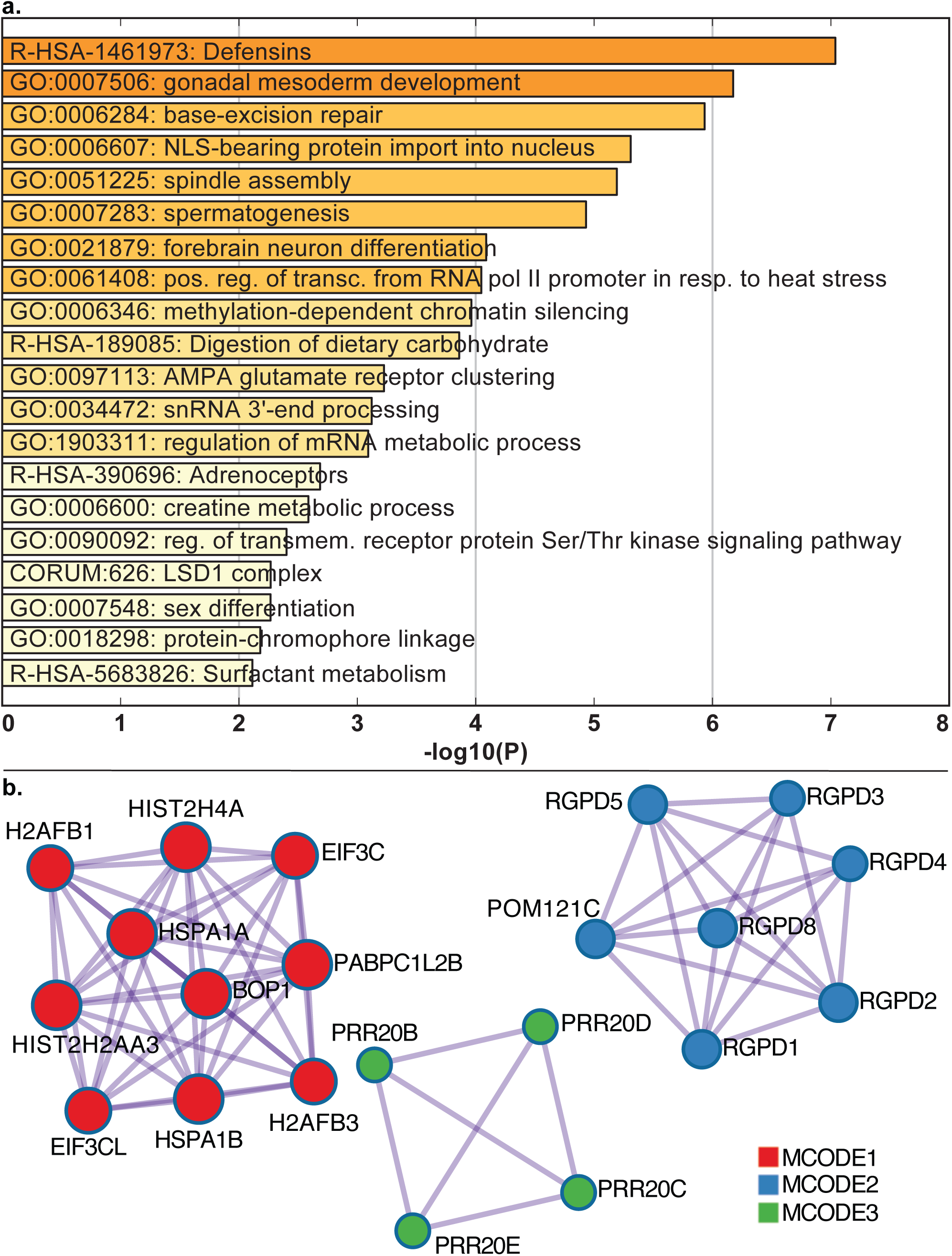
Pathways relevant to human health, development, and reproductive function are affected by dark and camouflaged genes. We characterized the pathways for dark and camouflaged genes using Metascape.org, including only genes where at least 5% of the CDS regions were dark (670 unique gene symbols; based on standard lllumina 100 nucleotide read lengths), **(a)** We identified several pathways that are important in human health, development, and reproductive function (Supplemental Table 11). Specific pathways included defensins (R-HSA-1461973; logP = −7.04), gonadal mesoderm development (G0:0007506; logP = −6.18), base-excision repair (G0:0006284; logP = −5.93), chromatin silencing (G0:0006342; logP = −5.86), Deubiquitination (R-HSA-5688426; logP = −5.32), NLS-bearing protein import into nucleus (G0:0006607; logP = −5.31), spindle assembly (G0:0051225; logP = −5.19), spermatogenesis (G0:0007283; logP = −4.93), and forebrain neuron differentiation (G0:0021879; logP = −4.09). **(b)** Looking specifically at known protein-protein interactions, Metascape identified 138 proteins with 212 known interactions (Supplemental Figure 4), and within those, identified three groups enriched for protein-protein interactions using the MCODE algorithm. All three MCODE groups combined are primarily associated with RNA transport (hsa030313; logP = −17.3; Supplemental Figure 5). Individually, the first group (MCODE1) is enriched for proteins involved in systemic lupus erythematosus (hsa05322; logP = −6.7), cellular response to stress (R-HSA-2262752; logP = −6.6), and RNA transport (hsa03013; logP = −4.39; Supplemental Figure 6). The second group (MCODE2) is enriched with proteins involved in NLS-bearing protein import into nucleus (G0:0006607; logP = −17.1) and protein import into nucleus (G0:0006606; logP = −15.4; Supplemental Figure 7). The third group does not have significant enrichment associations, likely because little is known about them; all four (*PRR20B, PRR20C, PRR20D*, and *PRR20E*) are 100% camouflaged and do not even have known expression measurements in GTEx [29] (Supplemental Figures 8-11).

Looking specifically at known protein-protein interactions, we found 138 proteins with 212 known interactions (Supplemental Figure 4), and within those, identified three groups enriched for protein-protein interactions using the MCODE algorithm [28] (Figure 4b). All three MCODE groups combined are primarily associated with RNA transport (hsa030313; logP = −17.3; Supplemental Figure 5; accessed December 2018). Individually, the first group (MCODE1) is enriched for proteins involved in systemic lupus erythematosus (hsa05322; logP = −6.7), cellular response to stress (R-HSA-2262752; logP = −6.6), and RNA transport (hsa03013; logP = −4.39; Supplemental Figure 6). The second group (MCODE2) is enriched with proteins involved in NLS-bearing protein import into nucleus (G0:0006607; logP = −17.1) and protein import into nucleus (G0:0006606; logP = −15.4; Supplemental Figure 7). The third group does not have significant enrichment associations, likely because little is known about them; all four genes (*PRR20B, PRR20C, PRR20D*, and *PRR20E*) are 100% camouflaged and do not even have known expression measurements in GTEx [29] (Supplemental Figures 8-11).

**Figure 6.**
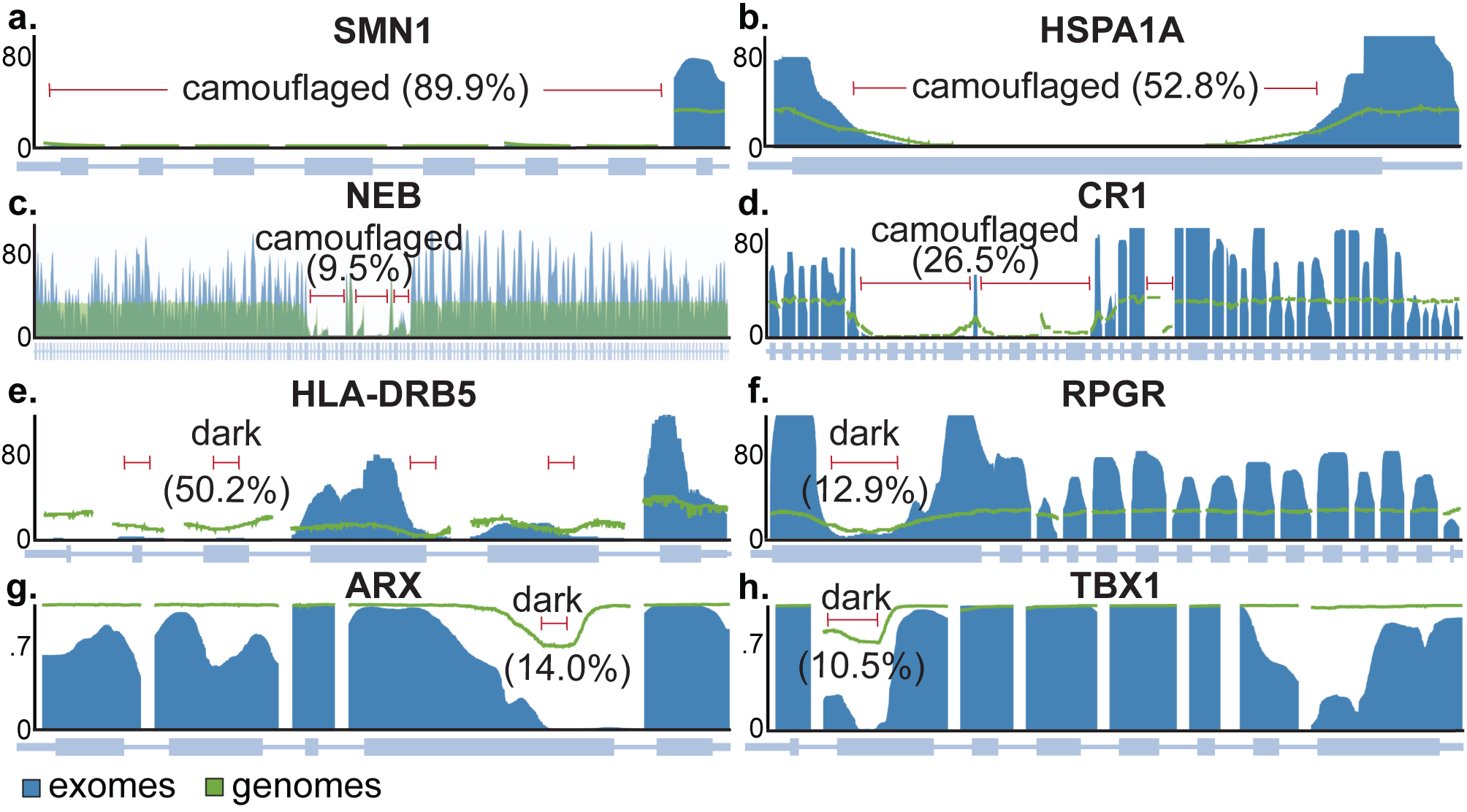
Camouflaged genes are consistently dark in gnomAD, but dark-by-depth genes may be sample or dataset specific. Most dark genes are specifically camouflaged (Supplemental Tables 12-13), but many are dark by depth; we found that camouflaged regions in the ADSP are consistently dark in the gnomAD consortium data (http://gnomad.broadinstitute.org/) [33]. Dark-by-depth regions may be more variable between samples and datasets, however, suggesting these regions may be sensitive to specific aspects of whole-genome sequencing (e.g., library preparation) or downstream analyses, **(a)** *SMN1* and *SMN2* are camouflaged by each other (89.9% and 88.2% dark CDS, respectively; only *SMN1* shown). Both genes contribute to spinal muscular atrophy, and have been implicated in ALS. **(b)** *HSPA1A* and *HSPA1B* are also camouflaged by each other (52.8% and 51.1% dark CDS, respectively; only *HSPA1A* shown). The heat-shock protein family has been implicated in ALS. **(c)** *NEB* (9.5% dark CDS) is a special case that is camouflaged by itself. *NEB* is associated with 24 diseases in the HGMD, including nemaline myopathy, a hereditary neuromuscular disorder. *NEB* is a large gene, thus, 9.5% dark CDS translates to 2424 protein-coding bases, **(d)** *CR1* is a top Alzheimer’s disease gene that plays a critical role in the complement cascade as a receptor for the C3b and C4b complement components, and potentially helps clear amyloid-beta (Aβ) [39-41]. *CR1* is also camouflaged by itself (26.5% dark CDS), where the repeated region includes the extracellular C3b and C4b binding domain. The number of repeats and density of certain isoforms have been associated with Alzheimer’s disease [21, 42-45]. **(e)** *HLA-DRB5* is dark by depth in the ADSP and gnomAD data (50.2% dark CDS). *HLA-DRB5* has been implicated in several diseases, including Alzheimer’s disease, **(f)** *RPGR* is likewise dark in ADSP and gnomAD (12.9% dark CDS), and is associated with several eye diseases, including retinitis pigmentosa and cone-rod dystrophy, **(g)** *ARX* is dark-by-depth (14.0% dark CDS), but varies by sample or cohort, as approximately 70% of gnomAD samples are not strictly dark by depth. *ARX* is associated with diseases including early infantile epileptic encephalopathy 1 (EIEE1) and Partington syndrome, **(h)** Similarly, *TBX1* is not strictly dark by depth in approximately 70% of gnomAD samples (10.5% dark CDS). The Y axes for figures **a-f** indicate median coverage in gnomAD (blue = exomes; green = genomes), whereas the Y axes in **g-h** represent the proportion of gnomAD samples that have >5x coverage. Dark and camouflaged regions, as well as the percentage of each gene’s CDS region that is dark, are indicated by red lines. Dark regions in exome data are either similar or more pronounced than what we observed in whole-genome data, highlighting that dark and camouflaged regions are generally magnified in whole-exome data. For interest, we also discovered that *APOE*—the top genetic risk for Alzheimer’s disease [34-36]—is approximately 6% dark CDS (by depth) for certain ADSP samples with whole-genome sequencing, and the same region is dark in gnomAD whole-exome data (Supplemental Figure 13).

**Figure 7.**
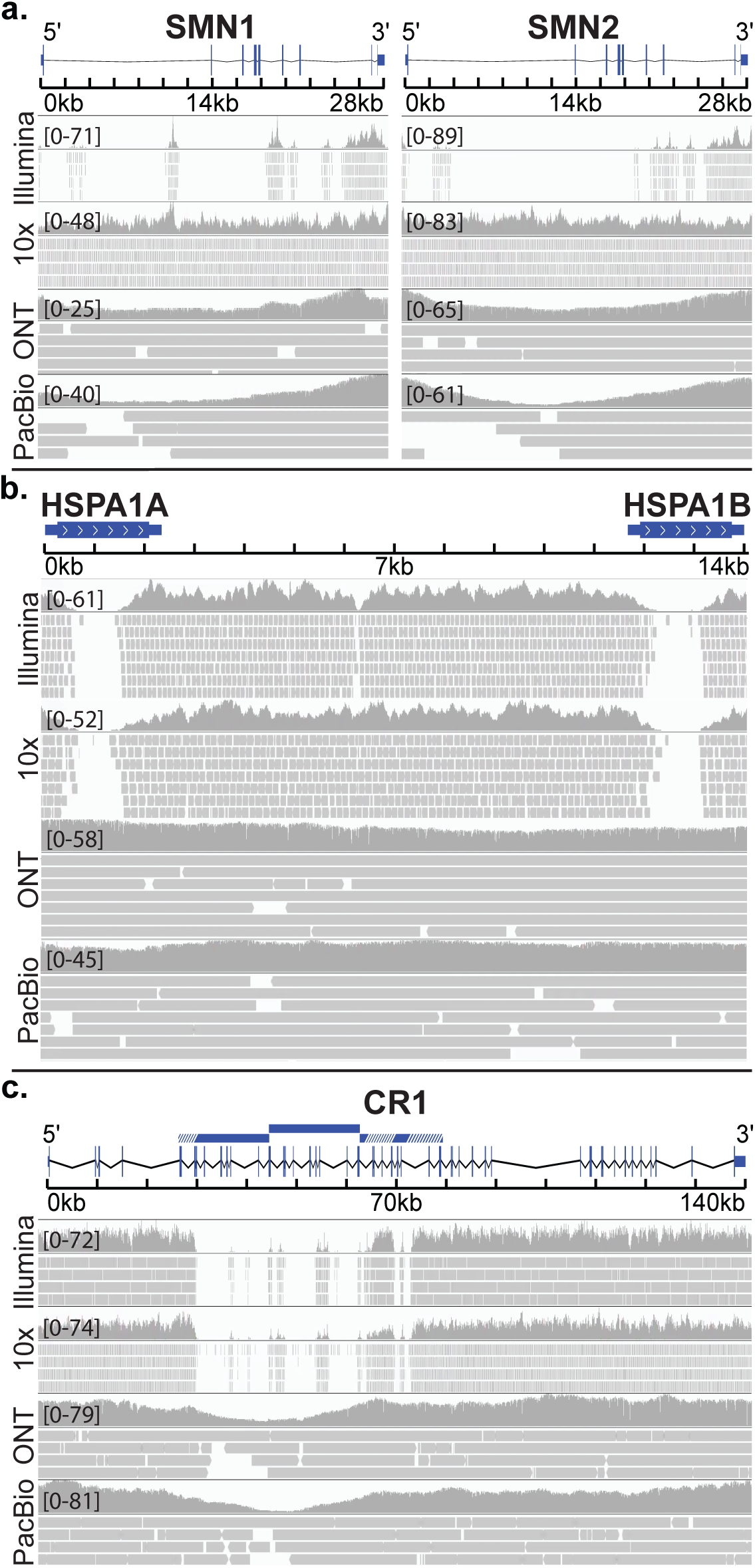
Long-read technologies resolve many camouflaged regions, with variable success. We found that ONT’s long-read technology appeared to resolve all camouflaged regions well with the high sequencing depth. PacBio performed similarly well, and lOx Genomics performs well under certain circumstances, **(a)** *SMN1* and *SMN2* were 89.9% and 88.2% dark CDS, respectively, based on standard lllumina sequencing with 100-nucleotide read lengths (illuminaRLlOO), and were 84.0% and 83.1% dark CDS based on lllumina 250-nucleotide read lengths (illuminaRL250; not shown). Both genes were technically 0% dark CDS for lOx Genomics, PacBio, and ONT data, **(b)** *HSPA1A* and *HSPA1B* were 52.8% and 51.1% dark CDS, respectively, based on illuminaRLlOO data, and were 50.2% and 49.5% dark CDS based on illuminaRL250 (not shown). Both genes were 0% dark CDS based on ONT and PacBio data, and were 45.8% and 51.8% dark CDS based on lOx Genomics data. In contrast to the results for *SMN1* and *SMN2*, both ONT and PacBio had consistent coverage throughout the camouflaged regions, whereas the camouflaged regions remain dark for lOx Genomics, **(c)** *CR1* was 26.5% dark CDS based on illuminaRLlOO, and was 24.5% dark based on illuminaRL250 (not shown). lOx Genomics did not improve coverage for *CR1*; the region remained 26.2% dark CDS. Both ONT and PacBio were 0% dark CDS. While both PacBio and ONT were able fill the camouflaged region, coverage dropped dramatically throughout the region, despite both genomes being sequenced at 50x and 46x median depth, which does not presently represent average use case for these technologies. The duplicated region is indicated by blue bars, where white lines indicate regions that have diverged sufficiently for reads to align uniquely. It is likely that the performance for ONT and PacBio long-read platforms will be better with longer sequencing libraries (e.g. >50kb fragment sizes). Regions were visualized with IGV. Reads with a MAPQ< 10 were filtered, and insertions, deletions, and mismatches are not shown.

**Figure 8.**
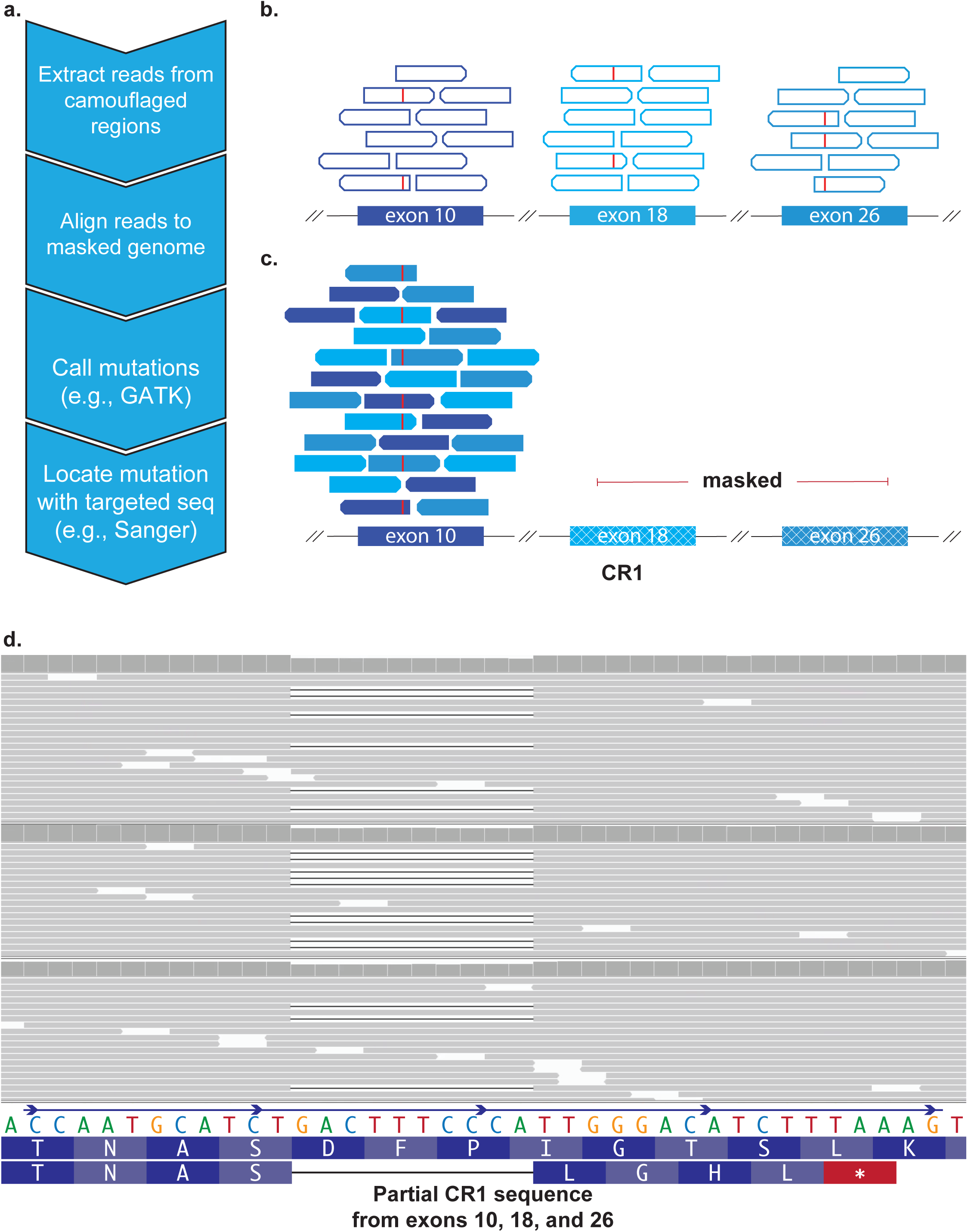
Many camouflaged regions can be rescued, including CR1, even in standard short-read sequencing data. Many large-scale whole-genome or whole-exome sequencing projects exist, covering tens of thousands of individuals. All of these datasets are affected by dark and camouflaged regions that may harbor mutations that either drive or modify disease in patients. Ideally, all samples would be re-sequenced using the latest technologies over time, but financial and biological samples are limited, making it essential to maximize the utility of existing data. We developed a method to rescue mutations in most camouflaged regions, including for standard short-read sequencing data. When confronted with a sequencing read that aligns to two or more regions equally well (with high confidence), most aligners (e.g., BWA [11-13]) will randomly assign the read to one of the regions with a low mapping quality (e.g., MAPQ = 0 for BWA). **(a)** Because the reads are already aligned to one of the regions, we can use the following steps to rescue mutations in most camouflaged regions: (1) extract reads from camouflaged regions; (2) mask all highly similar regions in the reference genome, except one, and re-align the extracted reads; (3) call mutations using standard methods (adjusting for ploidy); and (4) determine precise location using targeted sequencing (e.g., long-range PCR combined with Sanger, or targeted long-read sequencing [1]). Without competing camouflaged regions to confuse the aligner, the aligner will assign a high mapping quality, allowing variant callers to behave normally, **(b)** Exons 10,18, and 26 in *CR1* are identical, according to the reference genome. Standard aligners will randomly scatter reads matching that sequence across these exons and assign a low mapping quality (e.g., MAPQ = 0 for BWA; indicated as hollow reads). Red lines indicate an individual’s mutation that exists in one of these exons, but reads containing this mutation also get scattered and assigned a low mapping quality, **(c)** By masking exons 18 and 26, we can align all of these reads to exon 10 with high mapping qualities to determine whether a mutation exists. We cannot determine at this stage which of the three exons the mutation is actually located in, but researchers can test association with a given disease to determine whether the mutation is worth further investigation, **(d)** As a proof of principle, we rescued approximately 4622 exonic variants in the ADSP (TiTv = 1.97) using our method, including a frameshift mutation in *CR1* (MAF = 0.00019) that is only found in five cases and zero controls (three representative samples shown). The frameshift results in a stop codon shortly downstream. The ADSP is not large enough to formally assess association between the *CR1* frameshift and Alzheimer’s disease, but we believe the mutation merits follow-up studies given its location (*CR1* binding domain) and *CR1*’s strong association with disease.

**Figure 5.**
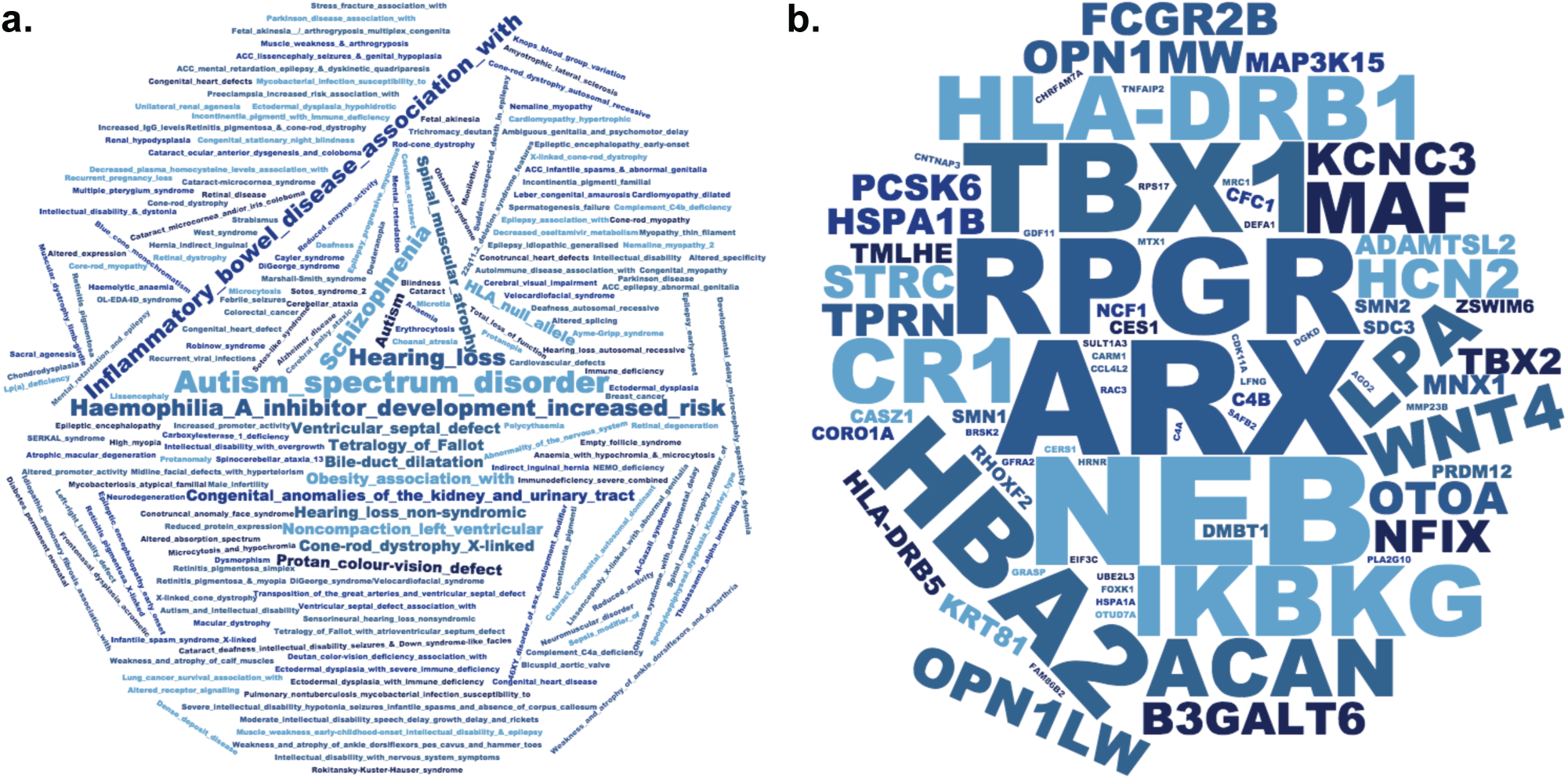
Seventy-five dark genes (≥5% CDS) are associated with 305 human phenotypes, including autism, inflammatory bowel disease, and others. We found 75 genes **≥**5% dark CDS that harbor mutations associated with 305 unique human phenotypes, including 277 diseases, according to the Human Gene Mutation Database (HGMD). **(a)** Some of the diseases with the most known associated genes include autism spectrum disorder, hemophilia A, schizophrenia, hearing loss, spinal muscular atrophy, and inflammatory bowel disease. Word size represents the number of genes associated with each disease. Some of the diseases most represented in our data are not surprising, given the number of genes involved in the disease, but these data demonstrate the number of diseases impacted by genes that are at least 5% dark CDS, and how important it is to completely resolve dark regions. We also performed an enrichment analysis, where the diseases most enriched for dark genes included Hemophilia A, color blindness (protan colour vision defect), and X-linked cone-rod dystrophy (Supplemental Figure 12). **(b)** Similarly, we quantified the number of diseases each gene was associated with, and identified many disease-relevant genes with large portions of dark CDS regions that may harbor critical disease-modifying mutations that currently go undetected. Some of the genes with the most known disease associations include *ARX* (14.0% dark CDS), *NEB* (9.5% dark CDS), *TBX1* (10.5% dark CDS), *RPGR* (12.9% dark CDS), *HBA2* (12.8% dark CDS), and *CR1* (26.5% dark CDS). *CR1* is particularly notable for neuroscientists and Alzheimer’s disease geneticists, patients, and their caregivers, given that CR1 is a top-ten Alzheimer’s disease gene. Other notable genes include SMN1 (89.9% dark CDS) and SMN2 (88.2% dark CDS), which are known to harbor mutations (in camouflaged regions) that are involved in spinal muscular atrophy (SMA) [65, 66, 113]. HSPA1A (52.8% dark CDS) and HSPA1B (51.1% dark CDS) also encode two primary 70-kilodalton (kDa) heat-shock proteins. Heat-shock proteins have been implicated in ALS [31, 32].

### There are 75 genes with known mutations associated with 305 human phenotypes

To assess the potential impact missing mutations in dark genes may have on human disease genetics, we measured the number of dark genes with at least 5% dark CDS that have mutations known to be involved in human disease; we ca1culated the number of genes that are ≥5% dark CDS with a mutation in the Human Gene Mutation Database (HGMD) [30]. We found 75 genes associated with 305 unique human phenotypes, including 277 diseases (Figure 5a). Some of the diseases with the most known associated genes include autism spectrum disorder, hemophilia A, schizophrenia, hearing loss, spinal muscular atrophy, and inflammatory bowel disease. Some of the diseases most represented in our data are not surprising, given the number of genes involved in the disease, but these data demonstrate the number of diseases impacted by genes that are at least 5% dark CDS. We also performed an enrichment analysis, where the diseases most enriched for dark genes included Hemophilia A, color blindness (protan colour vision defect), and X-linked cone-rod dystrophy (Supplemental Figure 12).

Similarly, we quantified the number of diseases each gene was associated with (Figure 5b). We identified many disease-relevant genes with large portions of dark CDS regions that may harbor critical disease-modifying mutations that currently go undetected. Some of the genes with the most known disease associations include *ARX* (14.0% dark CDS), *NEB* (9.5% dark CDS), *TBX1* (10.5% dark CDS), *RPGR* (12.9% dark CDS), *HBA2* (12.8% dark CDS), and *CR1* (26.5% dark CDS). The *CR1* gene is particularly notable given that *CR1* is a top-ten Alzheimer’s disease gene. Other notable genes include *SMN1* (89.9% dark CDS) and *SMN2* (88.2% dark CDS), which are known to be involved in spinal muscular atrophy (SMA) and ALS. *HSPA1A* (52.8% dark CDS) and *HSPA1B* (51.1% dark CDS) also encode two primary 70-kilodalton (kDa) heat-shock proteins, a family of proteins that have been implicated in ALS [31, 32].

### Camouflaged genes are consistently dark in gnomAD, but dark-by-depth genes may be sample or dataset specific

Although most dark genes are specifically camouflaged (Supplemental Tables 12-13), many are dark by depth in the ADSP data; upon manual comparison between whole-genome sequencing data from the ten ADSP males and coverage plots from the gnomAD consortium dataset (http://gnomad.broadinstitute.org/) [33], we found that camouflaged regions in the ADSP males are consistently dark in the gnomAD data, demonstrating that these camouflaged regions are consistent across datasets. The dark-by-depth regions are more variable between samples and datasets, however, suggesting these regions may be sensitive to specific aspects of whole-genome sequencing (e.g., library preparation) or downstream analyses. Specific camouflaged genes include *SMN1* and *SMN2* (89.9% and 88.2% dark CDS, respectively; Figure 6a), *HSPA1A* and *HSPA1B* (52.8% and 51.1% dark CDS, respectively; Figure 6b), *NEB* (9.5% dark CDS; Figure 6c), and *CR1* (26.5% dark CDS; Figure 6d). Specific dark-by-depth genes include *HLA-DRB5* (50.2% dark CDS; Figure 6e), *RPGR* (12.9% dark CDS; Figure 6f), *ARX* (14.0% dark CDS; Figure 6g), and *TBX1* (10.5% dark CDS; Figure 6h). All four camouflaged genes are also dark in the gnomAD data. A manual inspection of our dark-by-depth gene list, however, suggests most are not completely dark in gnomAD, but vary by sample or dataset. Specifically, *HLA-DRB5* and *RPGR* in gnomAD appear to be consistent with the ADSP data; *ARX* and *TBX1*, however, only appear to be dark in a portion of the gnomAD samples, where about 30% of samples have ≤5 reads in their respectively defined dark regions (Note: our threshold for dark regions is <5 reads, but the gnomAD plots for *ARX* and *TBX1* are based on <5 reads). Dark regions (Figures 6a-h) are either similar or more pronounced in the gnomAD whole-exome data than what we observed in the whole-genome data, highlighting that dark and camouflaged regions are generally magnified in whole-exome data; this is likely because of differences in library preparation and shorter read lengths in exome data. For interest, we also found that *APOE*—the top genetic risk for Alzheimer’s disease [34-36]—is approximately 6% dark CDS (by depth) for certain ADSP samples with whole-genome sequencing, and the same region is dark in gnomAD whole-exome data (Supplemental Figure 13). It is possible some of the dark regions we identified in standard short-read whole-genome data are specific to the ADSP samples, but additional work can clarify this issue. In either case, *dark-by-depth regions* (Supplemental Tables 14-15) *should be interrogated within individual datasets, and perhaps for individual samples as a quality control measure.*

*SMN1* and *SMN2* are camouflaged by each other, where both genes are known to contribute to spinal muscular atrophy, and have been implicated in ALS. *HSPA1A* and *HSPA1B* are also camouflaged by each other, and the heat-shock protein family has been implicated in ALS [37, 38]. *NEB* is a special case that is camouflaged by itself (rather than another gene), and is associated with 24 diseases in the HGMD, including nemaline myopathy, a hereditary neuromuscular disorder. *NEB* is a large gene (249151 nucleotides; 25577 CDS nucleotides), thus, ∼9.5% dark CDS translates to 2424 dark protein-coding bases. *CR1* is a top Alzheimer’s disease gene that plays a critical role in the complement cascade as a receptor for the C3b and C4b complement components, and potentially helps clear amyloid-beta (Aß) [39-41]. Like *NEB, CR1* is also camouflaged by itself, where the repeated region actually includes the extracellular C3b and C4b binding domain. The number of repeats and density of certain isoforms have been associated with Alzheimer’s disease [21, 42-45].

We found *HLA-DRB5* is dark by depth in the ADSP and gnomAD data, and has been implicated in several diseases, including Alzheimer’s disease. *RPGR* is likewise dark in ADSP and gnomAD, and is associated with several eye diseases, including retinitis pigmentosa and cone-rod dystrophy. We identified *ARX* as a dark-by-depth gene, but this gene appears to vary by sample or cohort, as only approximately 30% of gnomAD samples are strictly dark by depth, using our cutoff of <5 reads. *ARX* is associated with diseases including early infantile epileptic encephalopathy 1 (EIEE1) [46] and Partington syndrome [47]. Similarly, *TBX1*, which harbors mutations that cause the same phenotype as 22q11.2 deletion syndrome [48], is dark by depth in only approximately 30% of gnomAD samples.

### Long-read technologies resolve many camouflaged regions, with variable success

We selected three camouflaged gene regions to highlight common strengths and differences for how well each long-read sequencing technology addresses the camouflaged region, including *SMN1* and *SMN2* (Figure 7a), *HSPA1A* and *HSPA1B* (Figure 7b), and *CR1* (Figure 7c). The *SMN1* and *SMN2* genes are camouflaged by each other (gene duplication), as are *HSPA1A* and *HSPA1B. CR1*, however, is a special case, where it is camouflaged by a repeated region within itself. Only ONT appeared to be capable of fully addressing the camouflaged region for all three genes. lOx Genomics also performed well under certain circumstances, such as *SMN1* and *SMN2* (regions where the duplication is >50kb away), but did not perform well for *HSPA1A* and *HSPA1B.* PacBio performed well for *CR1* and *HSPA1A/HSPA1B*, but did not perform as well as ONT in the *SMN1/SMN2* region.

*SMN1* and *SMN2* were 89.9% and 88.2% dark CDS, respectively (Figure 7a), based on standard lllumina sequencing with 100-nucleotide read lengths, and were 84.0% and 83.1% dark CDS based on lllumina 250-nucleotide read lengths (not shown). Both genes were technically 0% dark CDS based on lOx Genomics, PacBio, and ONT data (Figure 7a). PacBio coverage does drop significantly throughout both genes, however.

*HSPA1A* and *HSPA1B* were 52.8% and 51.1% dark CDS (Figure 7b), respectively, based on standard lllumina 100-nucleotide read lengths, and were 50.2% and 49.5% dark CDS based on Illumina 250-nucleotide read lengths (not shown). Both genes were 0% dark CDS based on ONT and PacBio data, and were 45.8% and 51.8% dark CDS based on lOx Genomics data (Figure 7b). In contrast to the results for *SMN1* and *SMN2*, both ONT and PacBio had consistent coverage throughout the camouflaged regions, whereas the camouflaged regions remained dark for lOx Genomics (Figure 7b).

*CR1* was 26.5% dark CDS based on Illumina 100-nucleotide read lengths (Figure 7c), and was 24.5% dark based on Illumina 250-nucleotide read lengths (not shown). *CR1* was 26.2% dark CDS for lOx Genomics, and 0% for both ONT and PacBio (Figure 7c). While both PacBio and ONT were able fill the camouflaged region, coverage drops dramatically throughout the region, despite both genomes being sequenced at 50x and 46x median depth, which does not presently represent average use case for these technologies. It is likely that the performance of these long-read platforms will be better with longer average sequencing libraries (e.g. >50kb fragment sizes).

### Many camouflaged regions can be rescued, including in standard short-read sequencing data

There are many large-scale whole-genome or whole-exome sequencing projects across tens of thousands of individuals that are either completed or underway for a variety of diseases, including cancer (e.g., The Cancer Genome Atlas; TCGA), autism spectrum disorder (e.g., The Autism Sequencing Consortium; ASC), Alzheimer’s disease (e.g., The Alzheimer’s Disease Sequencing Project; ADSP), Parkinson’s disease (e.g., The Parkinson’s Progression Markers Initiative; PPMI), and ALS (e.g., Target ALS and CReATe). All of these datasets are affected by dark and camouflaged regions that may harbor mutations that are either driving or modify disease in patients. Ideally, all samples would be re-sequenced using the latest technologies over time, but financial resources and biological samples are limited, making it essential to maximize the utility of existing data.

Using a strategy similar to that proposed by Robert and Watson [10], we have developed a method to rescue mutations in most camouflaged regions, including for standard Illumina short-read sequencing data. When confronted with a sequencing read that aligns to two or more regions equally well (with high confidence), most aligners (e.g., BWA [11-13]) will randomly assign the read to one of the regions and assign a low mapping quality (MAPQ = 0 for BWA, or MAPQ = 1 for novoalign). Because the reads are already aligned to one of the regions, we can use the following steps to rescue mutations in most camouflaged regions (Figure 8): (1) extract reads from camouflaged regions; (2) mask all highly similar regions in the reference genome, except one, and re-align the extracted reads; (3) call mutations using standard methods. Without competing camouflaged regions to confuse the aligner, the aligner will assign a high mapping quality, allowing variant callers to behave normally. This will enable researchers to identify mutations that exist in one of the camouflaged regions, but not which specific region (Figure 8). After rescuing these mutations, researchers can then perform association studies to determine whether any of the mutations may be implicated in disease, and follow up with targeted sequencing methods to determine the exact camouflage region a mutation lies in.

### Re-alignment rescues approximately 4622 exonic variants, including a rare ten-nucleotide frameshift deletion in *CR1*

As a proof of principle, we applied our method to the Alzheimer’s Disease Sequencing Project (ADSP) case-control data [49] to approximate the number of potential mutations our approach could rescue. The ADSP is a large sequencing project organized, in part, to identify functional mutations that influence Alzheimer’s disease development. Across 13142 samples from the ADSP, excluding all variants with a quality by depth (QD) <2.5, we were able to rescue approximately 4622 exonic variants with a transition-transversion ration (Ti/Tv) of 1.97 from 147 camouflaged region sets, that are spread across 501 camouflaged genes (Supplemental Figure 14; VCF will be provided to the ADSP). Using a more stringent QD (excluding variants with QD <5), we rescued 3152 variants with a Ti/Tv ratio of 2.17. We only included camouflaged regions from CDS exons for all genes, including those that are <5% dark CDS.

Because *CR1* is a top-10 Alzheimer’s disease gene, we then specifically interrogated it using our method (Figure 8) for any functional mutations that could be involved in Alzheimer’s disease, and identified a rare ten-nucleotide frameshift deletion that is only found in five cases and zero controls, all of which are heterozygous (Figure 8d). Thus, the estimated minor allele frequency for this mutation is 5 / (13142 * 2) = 0.00019, making it more rare than the *TREM2* R47H allele [50-52]. For interest, only one of the individuals carried a single *APOEEA* allele (e3/e4). The other four individuals were homozygous for *APOE&3* (e3/e3). We were able to determine that the frameshift deletion is in one of exons 10,18, or 26. Briefly, our method extracts all reads with a low mapping quality (MAPQ < 10) from all three exons, masks all but one of the camouflaged regions within each set of camouflaged regions, and aligns all reads from each set to only one of the regions (Figure 8). Without identical competing regions to confuse the aligner, the mapping qualities are high enough for a variant caller (e.g., GATK HaplotypeCaller) to identify whether a mutation exists. For example, reads harboring the ten-nucleotide frameshift mutation were originally randomly scattered across exons 10,18, and 26 from the original alignment (Figure 8). We masked exons 18 and 26, leaving exon 10 unmasked; this allowed reads from each of the three exons to align to only exon 10, so we could perform variant calling. We estimate a cohort of approximately 70000 cases and controls would have approximately 80% statistical power to formally assess this mutation’s involvement in Alzheimer’s disease, assuming a Relative Risk (RR) of 3.3, at an alpha of 0.0001. We provide the .bed files in GRCh37 and GRCh38, along with scripts that will enable researchers to perform similar analyses in any sequencing dataset at https://github.com/mebbert/Dark_and_Camouflaged_genes.

## Discussion

While researchers have known for years that dark regions exist in standard short-read sequencing data, little work has been done to characterize the breadth of the issue, and to develop possible solutions until more financially-feasible long-read sequencing options are available. Short-read sequencing is unable to adequately address camouflaged regions because the reads cannot fully span camouflaged regions to properly align homologous nucleotides. Long-read sequencing technologies, such as those from lOx Genomics (synthetic long reads), Oxford Nanopore Technologies (ONT), and Pacific Biosciences (PacBio) have the potential to address many camouflaged regions because these technologies have median read lengths measured in thousands of nucleotides, rather than only 100-300 nucleotides from standard short-read sequencing technologies (e.g., Illumina). Recent work has even demonstrated that mappable ONT reads can exceed two million nucleotides (e.g, 2272580) [53, 54], showing future potential for addressing large camouflaged regions.

In this study, we systematically characterized dark and camouflaged gene regions and proposed a method to address most camouflaged regions in long- or short-read sequencing data. Our solution is specifically applicable to camouflaged regions, not regions that are dark by depth, simply because there are no reads available in regions that are dark by depth. While our solution is conceptually simple, implementing the solution systematically was challenging because of many intricate details, including increased zygosity, and would ideally be integrated into the original alignment and variant-calling process. While the original implementation was challenging, we provide the resulting .bed files for both GRCh37 and GRCh38 that are necessary to rescue mutations from camouflaged regions in any human re-sequencing dataset (https://github.com/mebbert/Dark_and_Camouflaged_genes). We also provide all of our data and source code. The .bed files and source code should make implementing our method relatively straightforward for other groups. As a proof of concept, we were able to rescue approximately 4622 variants in the ADSP dataset from 147 sets of camouflaged gene regions, which are spread across 501 camouflaged genes. Included in these rescued mutations is a ten-nucleotide frameshift deletion in *CR1* found in five ADSP cases and zero controls.

The number of genes affected by dark and camouflaged regions was surprisingly high. We identified 37873 total dark regions across 5857 gene bodies, nearly 4000 of which were protein coding genes. Exactly 28751 of the dark regions were intronic and 2657 were in protein-coding exons (CDS). Others were in pseudogenes (1134) and lincRNAs (732). While most of the dark regions were non-coding (e.g., intronic), these regions may still harbor important mutations that drive or modify human diseases. For example, there are many examples of mutations in non-coding regions driving disease, including repeat expansions [1, 55-62], splice-site mutations (these may be intronic or exonic) [63-77], and regulatory mutations (e.g., UTR regions) [78-87]. There are also many lincRNAs associated with disease [88-97].

There are many patients with diseases known to be genetically inherited that remain genetically unexplained because the patients do not have any of the known mutations. Many of the genes we identified as being at least partially dark are known to be involved in numerous diseases, including Alzheimer’s disease, ALS, SMA, hemophilia A, autism spectrum disorder, schizophrenia, and others; functional mutations that modify disease likely lie in some of these dark and camouflaged regions. For example, *SMN1* and *SMN2* are mostly dark (camouflaged) and are known to harbor mutations that cause disease [63, 65-67]. *CR1* is another dark gene that is 26.5% dark CDS, being camouflaged to itself, and is strongly implicated in Alzheimer’s disease. In fact, the *CR1* camouflaged region includes the C3b and C4b protein binding sites, repeated several times. Interestingly, the *C4B* gene (encodes the C4b protein) is also 72.8% dark CDS (camouflaged) and may be involved in disease [98, 99]. We are confident that rescuing mutations from camouflaged regions will have a meaningful impact on disease research, and may explain some of the missing heritability of Alzheimer’s disease [18, 100-102] and other diseases.

A large number of gene bodies (494) were 100% dark, which means they are entirely overlooked in standard whole-exome, whole-genome, and RNA sequencing studies [10]. Additionally, more than 1500 gene bodies, or nearly 27%, were at least 25% dark and more than 2000 (34.9%) were at least 5% dark; of these, 628 protein-coding genes were at least 5% dark within CDS regions. Understanding what role these genes play in human health and disease will require being able to resolve them in DNA and RNA sequencing experiments.

A critical decision for future large-scale sequencing projects will be regarding which long-read technology is ideal to maximize the probability of identifying functional mutations driving disease. Unfortunately, the answer is not clear, as each technology has its pros and cons. Based on our results, the ONT platform performed best, overall, resolving 71.4% of dark gene-body regions. Current costs will likely be prohibitive for large studies, however. The lOx Genomics platform resolved 66.3% of dark gene-body regions, when compared to standard Illumina sequencing. PacBio resolved 49.0% of dark gene-body regions. Even increasing Illumina read lengths from 100 to 250 made a sizeable difference, overall, resolving 21.1% of dark gene-body regions. Both the PacBio and ONT data used in this study had shorter median read lengths than expected, suggesting both technologies can likely perform better than our estimates.

Focusing only on CDS regions, there were 2757 dark CDS regions across 744 protein-coding genes, based on Illumina 100-nucleotide read lengths. ONT outperformed other long-read technologies, resolving 81.8% of dark CDS regions. PacBio and lOx Genomics resolved 66.6% and 54.9%, respectively. We found that lOx Genomics performed well in the *SMN1* and *SMN2* genes (Figure 7), attaining consistently deep, high-quality coverage throughout. Both ONT and PacBio coverage declined in the interior regions of the genes. In other cases, such as *CR1* and *NEB*, lOx Genomics was unable to improve on standard Illumina sequencing, but PacBio and ONT were able to largely resolve the region—albeit requiring higher than normal sequencing depth. We believe that lOx Genomics can correct the issues we observed in *CR1* and *NEB*, by implementing a more sophisticated version of our method that also incorporates evidence from their synthetic long-read technology.

Whether each technology is able to reliably resolve dark and camouflaged regions is an important consideration for choosing the best long-read technology, but we should also consider how reliably each technology is able to resolve structural mutations. In a previous study, we tested how well ONT and PacBio are able to traverse challenging repeat expansions, and whether they are amenable to genetic discovery [1]. We found that both technologies are well-suited, but we have not assessed performance of the lOx Genomics platform across long repeat expansions.

The primary challenge with ONT and PacBio long-read sequencing is, of course, the high error rate, which can be overcome through deeper sequencing because errors in ONT and PacBio sequencing are mostly random [103,104]. Ultimately, we are confident that, as long-read error rates improve, and costs continue to decline, long-read technologies will be the preferred sequencing choice for large-scale sequencing projects, especially when considering structural mutations.

We identified dark and camouflaged regions in this study by averaging data across ten males with deep lllumina whole-genome sequencing, using 100-nucleotide read lengths. We assessed how well long-read sequencing technologies (PacBio, ONT, and 10X genomics) resolve these regions, but our measurements should only be considered estimates. While long-read sequencing technologies are becoming more common, we were unable to find more than one male individual for each long-read technology; we needed male samples to assess all chromosomes, including the Y chromosome. Additionally, the samples we used for each long-read technology were sequenced at a much higher depth than is currently typical for a re-sequencingeffort, which is likely over estimating the number of dark regions they resolve for the average use case. Our measurements should be a reasonable estimate of reality, however, and future analyses will be able to refine our estimates.

We used whole-genome sequencing to assess dark and camouflaged regions, but this problem is magnified in whole-exome data, which many large-scale sequencing studies are based on, either completely, or in part. Whole-exome data are typically generated using even shorter read lengths. They are also generally based on capture, which means certain exons are not fully represented. *APOE* is a prime example, where it is typically well-covered in whole-genome data, but a portion is dark in whole-exome data (Supplemental Figure 13). With *APOE* harboring the largest genetic risk factors for Alzheimer’s disease, it is important to properly characterize the entire gene.

In this study, we characterized dark and camouflaged gene bodies, and demonstrated several disease-relevant genes where a significant portion is dark in standard short-read sequencing data, including *SMN1* and *SMN2, CR1*, and sometimes even *APOE.* We also identified a rare ten-nucleotide frameshift deletion in *CR1* that is found only in five ADSP cases and zero controls, as a proof of principle (Figure 8d). Using our method (Figure 8), we were able to determine that the frameshift deletion is in one of exons 10,18, or 26. With *CR1* being a top Alzheimer’s disease gene without any known functional mutations, we believe it will be important to assess this mutation in a large cohort, to determine whether it plays a role in disease development and progression. We have also proposed a solution to address most camouflaged genes in sequencing data, and believe that our approach has the potential to identify functional mutations that are influencing development across a range of diseases, but are currently entirely overlooked by standard short-read sequencing approaches.

## Conclusion

There remain thousands of potentially important genomic regions that are overlooked with short-read sequencing, but are largely resolved by long-read technologies. While these regions represent only a small portion of the entire genome or exome, many of these regions are known to be important in human health and disease. Equally important, however, is that the impact of many other genes is entirely unknown because they are 100% dark. We presented a method that can resolve most camouflaged regions that we believe will help researchers identify mutations that are involved in disease. As a proof of principle, we rescued approximately 4622 variants in the ADSP dataset, including a ten-nucleotide frameshift mutation in *CR1.* While we cannot formally assess the *CR1* frameshift mutation in Alzheimer’s disease (insufficient sample-size), we believe it is worth investigating in a larger cohort. In the long-term, we believe long-read sequencing technologies will be the best solution for resolving dark and camouflaged regions.

## Methods

### Sample selection and preparation

To identify dark and camouflaged regions, and to assess how well other technologies address them, we selected samples from each technology and read length. All samples were aligned to hg19/GRCh37. To assess dark and camouflaged regions in standard Illumina sequencing with 100-nucleotide read lengths, we selected ten unrelated male control samples from the Alzheimer’s Disease Sequencing Project (ADSP) where deep whole-genome sequencing had been performed by randomly selecting one male from ten random families. All ten males were from either the “Health/Medical/Biomedical“ (HMB-IRB) or “Health/Medical/Biomedical“ for non-profit organizations (HMB-IRB-NPU) consent groups, indicated as groups C1 and C2 in the ADSP pedigree files (available through dbGAP). We selected samples from the ADSP because we required samples that met the following criteria: (1) had been sequenced using standard paired-end lllumina sequencing with 10023 nucleotide read lengths, (2) had been sequenced with a median depth >30x, and (3) were publicly available. Median genome-wide read depths ranged from 35.4x to 42.9x, with a median of 39.4x. Samples were prepared and sequenced as part of the ADSP [49]. These samples were aligned using BWA (v0.5.9). We could not find samples from the 1000 Genomes Project [24] that met these criteria; sequencing depths were either too shallow, or read lengths were too long or short. The ADSP sample IDs we used were: A-CUHS-CU000406, A-CUHS-CU002997, A-CUHS-CU000779, A-CUHS-CU000208, A-CUHS-CU001010, A-CUHS-CU002031, A-CUHS-CU002707, A-CUHS-CU003023, A-CUHS-CU003090, A-CUHS-CU003128.

To assess dark and camouflaged regions in samples sequenced using Illumina 250-nucleotide read lengths, we selected ten samples from the 1000 Genomes Project that had been sequenced with 250-nucleotide read lengths, and had a median depth >30x. All ten samples were aligned using BWA (v 0.7.5a-r428) [2, 11-13]. Median genome-wide read depths ranged from 39.3 to 52.6, with a median of 48.9x. Sample IDs for the lllumina 250-nucleotide read lengths were: NA20845 (ftp://ftp.1000genomes.ebi.ac.uk/vol1/ftp/phase3/data/NA20845/high_coverage_alignment/), HG01112, HG01583 (ftp://ftp.1000genomes.ebi.ac.uk/vol1/ftp/phase3/data/HG01112/high_coverage_alignment/, HG01051 (ftp://ftp.1000genomes.ebi.ac.uk/vol1/ftp/phase3/data/HG01583/high_coverage_alignment/, HG03742 (ftp://ftp.1000genomes.ebi.ac.uk/vol1/ftp/phase3/data/HG01051/high_coverage_alignment/">ftp://ftp.1000genomes.ebi.ac.uk/vol1/ftp/phase3/data/HG01051/high_coverage_alignment/, HG00096 (ftp://ftp.1000genomes.ebi.ac.uk/vol1/ftp/phase3/data/HG03742/high_coverage_alignment/, HG01565 (ftp://ftp.1000genomes.ebi.ac.uk/vol1/ftp/phase3/data/HG00096/high_coverage_alignment/, HG01879 (ftp://ftp.1000genomes.ebi.ac.uk/vol1/ftp/phase3/data/HG01565/high_coverage_alignment/, HG01500 (ftp://ftp.1000genomes.ebi.ac.uk/vol1/ftp/phase3/data/HG01879/high_coverage_alignment/, and HG03006 (ftp://ftp.1000genomes.ebi.ac.uk/vol1/ftp/phase3/data/HG01500/high_coverage_alignment/ (ftp://ftp.1000genomes.ebi.ac.uk/voll/ftp/phase3/data/HG03006/high_coverage_alignment/

We also selected samples generated using the lOx Genomics synthetic long-read sequencing platform, and ONT and PacBio long-read sequencing platforms that were either prepared by, and publicly available from the respective company, or prepared using standard practice. Specifically, we downloaded HG00512 raw FASTQ data from lOx Genomics (https://support.10xgenomics.eom/de-novo-assembly/datasets/l.l.0/msHG00512; http://s3-us-west-2.amazonaws.eom/10x.files/samples/assemblv/2.l.0/chi/chifastqs.tar) and aligned it according to lOx Genomics’ standard practices. We used longranger (v2.2.2) and aligned to GRCh37 (longranger wgs —id HG00512 — description=“Han Chinese“ -sex=“male” — fastqs=chi/HNKHFCCXX/,chi/HWHFTCCXX/ -reference=“10x-b37-2.1.0/“ -jobmode=sge -mempercore=125 - downsample=385). Median depth for HG00512 was 52x, after downsampling. For ONT, we downloaded the final Cliveome v2 from ONT’s official GitHub page (http://cliveo.me/: https://github.com/nanoporetech/ONT-HGl/blob/master/CONTENTS.md), which was prepared by ONT. Cliveome v2 was sequenced to a median depth of 36x. To increase the median read depth to more closely match those of other technologies, we merged reads from HG002 (https://ftp-trace.ncbi.nlm.nih.gov/giab/ftp/data/AshkenazimTrio/HG002_NA24385_son/Ultralong_OxfordNanopore/com_bined_2018-08-10/HG0020NTrel216xRGHPlOxtrioRTG.cram) [105,106] and aligned using minimap2 [107] (ALIGN_OPTS=“x map-pb -a -eqx -L -O 5,56 -E 4,1 -B 5 -secondary=no -z 400,50 -r 2k -Y“; REF=glkv37/glkv37.fa; minimap2 -d ${REF}.mmi ${ALIGN_OPTS} ${REF}; minimap2 ${ALIGN_OPTS} -a ${REF}.mmi <reads.fq> | samtools view -T {REF} -F 2308 > output_file). The merged sample had 46x median depth. We used the same alignment options recommended for PacBio because we found the recommended ‘map-ont’ option in minimap2 performed substantially worse. We used PacBio data generated from HG005 (ftp://ftp-trace.ncbi.nlm.nih.gov/giab/ftp/data/ChineseTrio/HG005_NA24631_son/MtSinai_PacBio/PacBio_minimap2_bam/) [105], which was sequenced to a median depth of 50x and aligned using minimap2 [107] (pbsv fasta [movie].subreads.bam | minimap2 -t 8 -x map-pb -a -eqx -L-0 5,56 -E 4,1 -B 5 -secondary=no -z 400,50-r 2k - ftp://ftp.1000genomes.ebi.ac.uk/vol1/ftp/technical/reference/phase2_reference_assembly_sequence/hs37d 5.<u>fa.gz</u> - | samtools sort > HG005_PacBio_GRCh37.bam). Neither the ONT nor the PacBio alignments include secondary alignments.

### Identifying dark and camouflaged gene body regions

To identify dark and camouflaged gene body regions in standard Illumina 100-nucleotide read length data, we first scanned all ten ADSP whole-genome sequence samples for genomic positions that met either of the following criteria: (1) had <5 reads, and (2) had ≥90% of reads with a mapping quality (MAPQ) <10. We then averaged the depth and count of low MAPQ reads across all samples for each position. We used strict cutoffs to identify regions that are clearly dark, but there are many additional regions that fall just beyond our thresholds. This analysis was performed using the Dark Region Finder (DRF; https://github.com/mebbert/DarkRegionFinder; mapq=9; dark_mass=90; camo_mass=50; dark_depth=5; java -jar -Xmx20g CamoGeneFinder.jar -i <sample>.bam -human-ref genome.fa -min-region-size 1 -camo-mapq-threshold $mapq —min-dark-mapq-mass $dark_mass —min-camo-mapq-mass $camo_mass -dark-depth $dark_depth -camo-bed-output <sample>-camo-dark_depth_${dark_depth}-dark_mass_${dark_mass}-camo_mass_${camo_mass}-mapq_${mapq}.b37.bed -dark-bed-output <sample>-dark-dark_depth_${dark_depth}-dark_mass_${dark_mass}.b37.bed -incomplete-bed-output <sample>- incomplete.b37.bed). Any position that met either criteria was considered dark and categorized as either dark by depth or dark by mapping quality. We then limited the dark regions to gene bodies by intersecting dark regions identified by Dark Region Finder with Ensembl’s GRCh37 build 87 gene annotations. We converted the transcript-level annotations to gene-level annotations using bedtools [108] and custom scripts that are available. Any dark region that spanned a gene body element region (e.g., intron-exon boundary) was split into two separate dark regions so we could estimate the number of dark bases in each type of gene body region (e.g., introns, exons, UTRs, etc.). For most analyses, we only included dark regions with ≥20 contiguous bases. The only exception is for Supplemental Tables 1, 3, 5, 7, 9,12, and 14, where we calculate total percentage of each gene body that is dark, in which we include all dark positions. To identify camouflaged regions, specifically, we used BLAT [26] to identify all genomic regions that were highly similar to any given gene body region that was dark by mapping quality. Any region that was ≥98% identical (-minIdentity = 98), and that was considered dark (≥90% of reads with MAPQclO), was considered a match. We generated .bed files for GRCh37 using this method. We also converted the GRCh37 .bed file to GRCh38 using a custom script, based off the Ensembl build 87 GRCh38 gene annotations. All code and .bed files can be found at https://github.com/mebbert/Dark_and_Camouflaged_genes.

### Statistics

We quantified the percentage of each gene body that was dark by summing the total number of dark bases in the gene (i.e., between the 5’UTR to the 3’UTR start and end, respectively) and dividing by the total number of bases in the gene. We similarly calculated the percentage of intronic, exonic (including CDS and UTR), and only CDS exons by dividing the total number of dark bases in each category within the gene by the total number of bases within that category. We performed these calculations for data based on Illumina 100-nucleotide reads for all dark regions combined (Supplemental Tables 1-2), dark by depth only (Supplemental Tables 14-15), dark by mapping quality (Supplemental Tables 16-17), and only camouflaged regions (Supplemental Tables 12-13). We performed identical calculations for the samples from Illumina 250-nucleotide read length data, lOx Genomics, ONT, and PacBio (Supplemental Tables 3-10,18-41). We identified diseases that were known to be associated with genes that are at least 5% dark CDS by searching for mutations in the Human Gene Mutation Database (HGMD) [30].

Coverage plots from gnomAD data were obtained from gnomAD-old.broadinstitute.org [33]. We used the old version because the current version of gnomAD (accessed December 2018) does not allow the user to view median read depths, nor the percentage of samples with greater than a given coverage depth. Sequence pileups in representative samples were generated using the Integrative Genomics Viewer (IGV) [109], where reads with a MAPQ < 10 were filtered, and insertions, deletions, and mismatches were not shown. Karyotype plots showing genomic locations for dark and camouflaged regions were generated using KaryotypeR (vl.6.2) [110] in R (v3.5.1). Bar plots were made using ggplot2 (v3.0.0). Pathway analyses and resulting plots were generated using Metascape (accessed December 2018) [111]. Word clouds were generated at wordclouds.com. Gene schematics were generated using the Gene Structure Display Server (GSDS; v2) [112].

We performed an enrichment analysis to assess whether genes that are ≥5% dark CDS are enriched for specific diseases. Because we identified 75 genes that have a known mutation associated with disease, and that are >5% dark CDS, we randomly selected 75 genes from the with known HGMD mutations and measured the number of genes with known mutation associated with each disease. We repeated this process 10000 times and used the following metric as our enrichment score: -10*logl0(empirical_pvalue), rounded to the nearest whole number.

### Screening ADSP for functional *CR1* mutations in camouflaged region

After discovering that more than 25% of the *CR1* gene’s CDS is camouflaged, we screened all ADSP samples for rare functional mutations that could play a role in Alzheimer’s disease development and progression by applying our proposed method (Figure 8). To apply our method, we extracted all reads with a mapping quality (MAPQ) <10 from each camouflaged region within *CR1*, and from each of the respective camouflage mate regions, using samtools and the GRCh37 .bed file we generated that identifies all camouflaged regions. An example of camouflaged mate regions in *CR1* includes exons 10,18, and 26, which are identical in the reference genome (Figure 8). As previously mentioned, *CR1* is a special case that is camouflaged by regions duplicated within itself, rather than being camouflaged by a different gene; thus, we knew that any mutations we discovered would be from *CR1.* Our approach works the same regardless of whether a gene is camouflaged by itself or another gene, but we mention that *CR1* is camouflaged by itself, for interest. After extracting reads from each camouflaged region, using the .bed file we provide, we then masked all camouflaged regions within *CR1* in the reference genome, except for one from each set of camouflaged mates. For example, between exons 10,18, and 26, we masked exons 18 and 26 in the reference genome, allowing reads from all three exons to align only to exon 10; without competing camouflaged regions to confuse the aligner, all reads from exons 10,18, and 26 mapped to exon 10 with high quality. Masking regions of the reference genome simply means to change nucleotides to an unmappable character (usually ‘N’), to prevent any reads from aligning to that region.

After aligning all reads to a single region within each set of camouflaged regions, we were able to perform standard variant calling using the GATK HaplotypeCaller [25], with one exception: instead of treating each camouflaged region as diploid, we increased the ploidy setting in HaplotypeCaller according to the number of copies within a given set of camouflaged regions. Referring again to our *CR1* example, because there are three regions (exons 10,18, and 26), we set the HaplotypeCaller ploidy to hexaploid. Increasing the ploidy is essential for increased sensitivity, since the number of reads harboring a given variant—which only originate from one of the camouflaged regions—will be overwhelmed by reads from the others, thus preventing the variant caller from identifying the mutation under the assumption that the data are from a diploid region. In other words, if a mutation exists in exon 26, we would expect only approximately l/6^th^ of reads from exons 10,18, and 26 to harbor that mutation, rather than approximately 1/2. Because the ADSP is mostly exome data, we limited HaplotypeCaller to CDS exons only. According to the current ADSP phenotype data, one of the samples harboring the *CR1* frameshift mutation is a control. The individual has since been officially diagnosed with Alzheimer’s disease, however.

## Abbreviations

MAPQ: mapping quality; CDS: coding sequence; ALS: amyotrophic lateral sclerosis; FTD: frontotemporal dementia; PacBio: Pacific Biosciences; ONT: Oxford Nanopore Technologies; ADSP: Alzheimer’s Disease Sequencing Project;

## Declarations

### Ethics approval and consent to participate

The Mayo Clinic Institutional Review Board (IRB) approved all procedures for this study and we followed all appropriate protocols.

### Consent for publication

All participants were properly consented for this study.

### Availability of data and materials

The ADSP dataset supporting the conclusions of this article (including the whole-genome and whole-exome data) are available in the National Institute on Aging Genetics of Alzheimer’s Disease Storage (NIAGADS) site, and may be requested therein: https://www.niagads.org/adsp/. Public links to the high-coverage whole-genome data from the 1000 Genomes Project (illumina 250bp read lengths) are listed in the Methods. Raw data from the 10x Genomics sample (HG00512) used within this article was downloaded directly from the 10x Genomics website at: https://support.10xgenomics.eom/de-novo-assemblv/datasets/2.l.0/chi. The Cliveome2 data was downloaded from the official Oxford Nanopore Technologies GitHub page: https://github.com/nanoporetech/ONT-HGl/blob/master/. The PacBio data used in this publication (HG005) was downloaded from ftp://ftp-trace.ncbi.nlm.nih.gov/giab/ftp/data/ChineseTrio/HG005_NA24631_son/MtSinai_PacBio/PacBio_minimap2_bam/HG005_PacBio_GRCh37.bam.

All scripts are available at: https://github.com/mebbert/Dark_and_Camouflaged_genes.

### Competing interests

All authors declare they have no conflicts of interest.

### Funding

This work was supported by the PhRMA Foundation [RSGTMT17 to M.E.]; the Ed and Ethel Moore Alzheimer’s Disease Research Program of Florida Department of Health [8 AZ10 and 9AZ08 to M.E., and 6AZ06 to J.F.]; the Muscular Dystrophy Association (M.E.); the National Institutes of Health [NS094137 to J.F., AG047327 to J.F, AG049992 to J.F., NS097261 to R.R., NS097273 to L.P., NS084528 to L.P., NS084974 to L.P., NS099114 to L.P., NS088689 to L.P., NS093865 to L.P.]; Department of Defense [ALSRP AL130125 to L.P.]; Mayo Clinic Foundation (L.P. and J.F.); Mayo Clinic Center for Individualized Medicine (L.P. and J.F.); Amyotrophic Lateral Sclerosis Association (M.E., L.P.); Robert Packard Center for ALS Research at Johns Hopkins (L.P.) Target ALS (L.P.); Association for Frontotemporal Degeneration (L.P.); GHR Foundation (J.F.); and the Mayo Clinic Gerstner Family Career Development Award (J.F.).

### Authors’ contributions

ME, LP, and JF developed and designed the study, and wrote the manuscript. ME and TJ performed all analyses. JR, SY, NT, YA, VB, EL, DK, PC, LP, PR, JK, MC, OA, and RR contributed important intellectual ideas and feedback. SY, YA, NT, OA, and RR helped obtain data. KW and JS performed experiments. EL, DK, and PC provided samples. All authors read and approved the final manuscript.

## Acknowledgements

Biological samples and Associated Phenotypic Data used in primary data analyses were stored at Principal Investigators’ institutions, and at the National Cell Repository for Alzheimer’s Disease (NCRAD) at Indiana University funded by NIA. Associated Phenotypic Data used in primary and secondary data analyses were provided by Principal Investigators, the NIA funded Alzheimer’s Disease Centers (ADCs), and the National Alzheimer’s Coordinating Center (NACC) and stored at Principal Investigators’ institutions, NCRAD, and at the National Institute on Aging Alzheimer’s Disease Data Storage Site (NIAGADS) at the University of Pennsylvania, funded by NIA. Contributors to the Genetic Analysis Data included Principal Investigators on projects that were individually funded by NIA, other NIH institutes, private U.S. organizations, or foreign governmental or nongovernmental organizations.

The Alzheimer’s Disease Sequencing Project (ADSP) is comprised of two Alzheimer’s Disease (AD) genetics consortia and three National Human Genome Research Institute (NHGRI) funded Large Scale Sequencing and Analysis Centers (LSAC). The two AD genetics consortia are the Alzheimer’s Disease Genetics Consortium (ADGC) funded by NIA (U01AG032984), and the Cohorts for Heart and Aging Research in Genomic Epidemiology (CHARGE) funded by NIA (R01 AG033193), the National Heart, Lung, and Blood Institute (NHLBI), other National Institute of Health (NIH) institutes and other foreign governmental and non-governmental organizations. The Discovery Phase analysis of sequence data is supported through UF1AG047133 (to Drs. Schellenberg, Farrer, Pericak-Vance, Mayeux, and Haines); U01AG049505 to Dr. Seshadri; U01AG049506 to Dr. Boerwinkle; U01AG049507 to Dr. Wijsman; and U01AG049508 to Dr. Goate and the Discovery Extension Phase analysis is supported through U01AG052411 to Dr. Goate, U01AG052410 to Dr. Pericak-Vance and U01 AG052409 to Drs. Seshadri and Fornage. Data generation and harmonization in the Follow-up Phases is supported by U54AG052427 (to Drs. Schellenberg and Wang).

The ADGC cohorts include: Adult Changes in Thought (ACT), the Alzheimer’s Disease Centers (ADC), the Chicago Health and Aging Project (CHAP), the Memory and Aging Project (MAP), Mayo Clinic (MAYO), Mayo Parkinson’s Disease controls, University of Miami, the Multi-Institutional Research in Alzheimer’s Genetic Epidemiology Study (MIRAGE), the National Cell Repository for Alzheimer’s Disease (NCRAD), the National Institute on Aging Late Onset Alzheimer’s Disease Family Study (NIA-LOAD), the Religious Orders Study (ROS), the Texas Alzheimer’s Research and Care Consortium (TARC), Vanderbilt University/Case Western Reserve University (VAN/CWRU), the Washington Heights-Inwood Columbia Aging Project (WHICAP) and the Washington University Sequencing Project (WUSP), the Columbia University Hispanic-Estudio Familiar de Influencia Genetica de Alzheimer (EFIGA), the University of Toronto (UT), and Genetic Differences (GD).

The CHARGE cohorts are supported in part by National Heart, Lung, and Blood Institute (NHLBI) infrastructure grant HL105756 (Psaty), RC2HL102419 (Boerwinkle) and the neurology working group is supported by the National Institute on Aging (NIA) R01 grant AG033193. The CHARGE cohorts participating in the ADSP include the following: Austrian Stroke Prevention Study (ASPS), ASPS-Family study, and the Prospective Dementia Registry-Austria (ASPS/PRODEM-Aus), the Atherosclerosis Risk in Communities (ARIC) Study, the Cardiovascular Health Study (CHS), the Erasmus Rucphen Family Study (ERF), the Framingham Heart Study (FHS), and the Rotterdam Study (RS). ASPS is funded by the Austrian Science Fond (FWF) grant number P20545-P05 and P13180 and the Medical University of Graz. The ASPS-Fam is funded by the Austrian Science Fund (FWF) project 1904), the EU Joint Programme - Neurodegenerative Disease Research (JPND) in frame of the BRIDGET project (Austria, Ministry of Science) and the Medical University of Graz and the Steiermärkische Krankenanstalten Gesellschaft. PRODEM-Austria is supported by the Austrian Research Promotion agency (FFG) (Project No. 827462) and by the Austrian National Bank (Anniversary Fund, project 15435. ARIC research is carried out as a collaborative study supported by NHLBI contracts (HHSN268201100005C, HHSN268201100006C, HHSN268201100007C, HHSN268201100008C, HHSN268201100009C, HHSN268201100010C, HHSN268201100011C, and HHSN268201100012C). Neurocognitive data in ARIC is collected by U01 2U01HL096812, 2U01HL096814, 2U01HL096899, 2U01HL096902, 2U01HL096917 from the NIH (NHLBI, NINDS, NIA and NIDCD), and with previous brain MRI examinations funded by R01-HL70825 from the NHLBI. CHS research was supported by contracts HHSN268201200036C, HHSN268200800007C, N01HC55222, N01HC85079, N01HC85080, N01HC85081, N01HC85082, N01HC85083, N01HC85086, and grants U01HL080295 and U01HL130114 from the NHLBI with additional contribution from the National Institute of Neurological Disorders and Stroke (NINDS). Additional support was provided by R01AG023629, R01AG15928, and R01AG20098 from the NIA. FHS research is supported by NHLBI contracts N01-HC-25195 and HHSN268201500001I. This study was also supported by additional grants from the NIA (ROls AG054076, AG049607 and AG033040 and NINDS (R01 NS017950). The ERF study as a part of EUROSPAN (European Special Populations Research Network) was supported by European Commission FP6 STRP grant number 018947 (LSHG-CT-2006-01947) and also received funding from the European Community’s Seventh Framework Programme (FP7/2007-2013)/grant agreement HEALTH-F4-2007-201413 by the European Commission under the programme “Quality of Life and Management of the Living Resources” of 5th Framework Programme (no. QLG2-CT-2002-01254). High-throughput analysis of the ERF data was supported by a joint grant from the Netherlands Organization for Scientific Research and the Russian Foundation for Basic Research (NWO-RFBR 047.017.043). The Rotterdam Study is funded by Erasmus Medical Center and Erasmus University, Rotterdam, the Netherlands Organization for Health Research and Development (ZonMw), the Research Institute for Diseases in the Elderly (RIDE), the Ministry of Education, Culture and Science, the Ministry for Health, Welfare and Sports, the European Commission (DG XII), and the municipality of Rotterdam. Genetic data sets are also supported by the Netherlands Organization of Scientific Research NWO Investments (175.010.2005.011, 911-03-012), the Genetic Laboratory of the Department of Internal Medicine, Erasmus MC, the Research Institute for Diseases in the Elderly (014-93-015; RIDE2), and the Netherlands Genomics Initiative (NGI)/Netherlands Organization for Scientific Research (NWO) Netherlands Consortium for Healthy Aging (NCHA), project 050-060-810. All studies are grateful to their participants, faculty and staff. The content of these manuscripts is solely the responsibility of the authors and does not necessarily represent the official views of the National Institutes of Health or the U.S. Department of Health and Human Services.

The four LSACs are: the Human Genome Sequencing Center at the Baylor College of Medicine (U54 HG003273), the Broad Institute Genome Center (U54HG003067), The American Genome Center at the Uniformed Services University of the Health Sciences (U01AG057659), and the Washington University Genome Institute (U54HG003079).

Biological samples and associated phenotypic data used in primary data analyses were stored at Study Investigators institutions, and at the National Cell Repository for Alzheimer’s Disease (NCRAD, U24AG021886) at Indiana University funded by NIA. Associated Phenotypic Data used in primary and secondary data analyses were provided by Study Investigators, the NIA funded Alzheimer’s Disease Centers (ADCs), and the National Alzheimer’s Coordinating Center (NACC, U01AG016976) and the National Institute on Aging Genetics of Alzheimer’s Disease Data Storage Site (NIAGADS, U24AG041689) at the University of Pennsylvania, funded by NIA, and at the Database for Genotypes and Phenotypes (dbGaP) funded by NIH. This research was supported in part by the Intramural Research Program of the National Institutes of health, National Library of Medicine. Contributors to the Genetic Analysis Data included Study Investigators on projects that were individually funded by NIA, and other NIH institutes, and by private U.S. organizations, or foreign governmental or nongovernmental organizations.

