## Supplemental figures for "Systematic analysis of dark and camouflaged genes: disease-relevant genes hiding in plain sight"

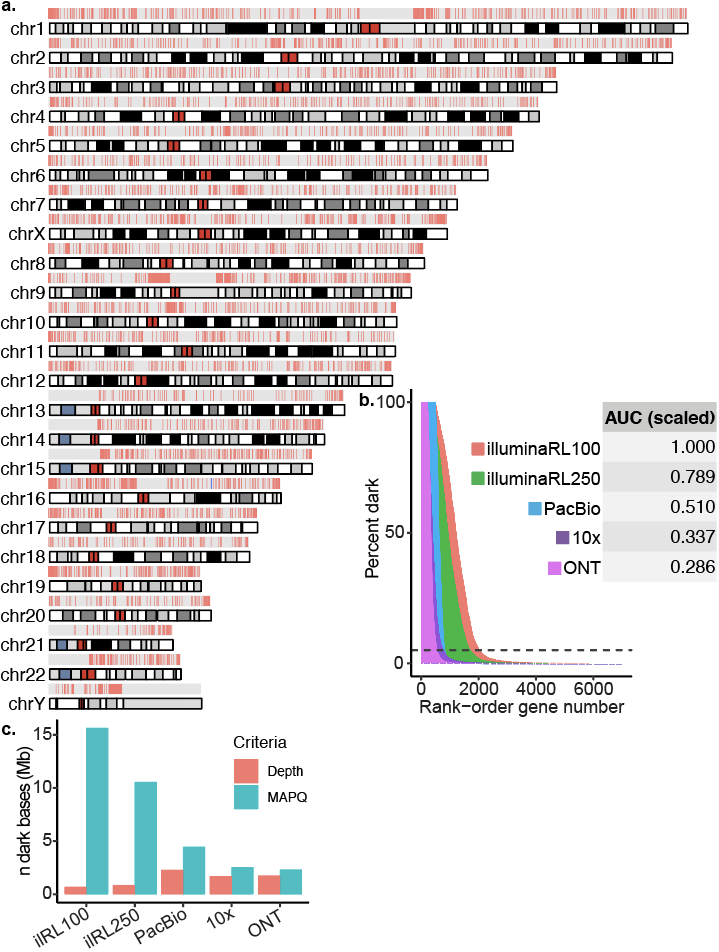

**Supplemental Figure 1. Dark regions are scattered throughout the genome, but largely resolved by long-read sequencing technologies.** **(a)** We identified 37873 dark regions (>16 million nucleotides) in 5857 gene bodies (based on Ensemble GRCh37 build 87 gene annotations) that were either dark by depth or dark by mapping quality (Supplemental Tables 1-2). Stratifying the gene bodies by GENCODE biotype [27], 3635 gene bodies were protein coding, 1102 were pseudogenes, and 720 were long intergenic non-coding RNAs (lincRNA) (Figure 2a). Of all 37873 dark gene-body regions, 28598 were intronic, 4113 were in non-coding RNA exons (e.g., lincRNAs and pseudogenes), 2657 were in protein-coding exons (CDS), 1134 were in 3’UTR regions, and 1103 were in 5’UTR regions (Figure 2b; Supplemental Table 1). Any dark region that spanned a gene element boundary (e.g., intron to exon) was split into separate dark regions. Of the 5857 gene bodies, 494 (8.4%) were 100% dark, 1560 (26.6%) were at least 25% dark, and 2046 (34.9%) were at least 5% dark Supplemental Table 1). **(b)** Data from samples sequenced using 250-nucleotide Illumina read lengths reduced the area under the curve by 21.1% for all gene bodies (Supplemental Tables 3-4); this translates to reducing the number of dark nucleotides by 30.1%. Comparing long-read sequencing technologies to the standard Illumina 100-nucleotide read lengths, PacBio, 10x Genomics, and ONT reduced the area under the curve (AUC) by approximately 49.0%, 66.3%, and 71.4% for all gene bodies, respectively (Supplemental Tables 5-10); this translates to reducing the number of dark nucleotides by 58.8%, 74.2%, and 75.1%, respectively. The ONT platform performed best, overall, reducing the total percentage of dark regions by 71.4% to 28.6%. **(c)** Long-read technologies improve upon short-read data mostly by reducing the percentage of regions that are dark by mapping quality.

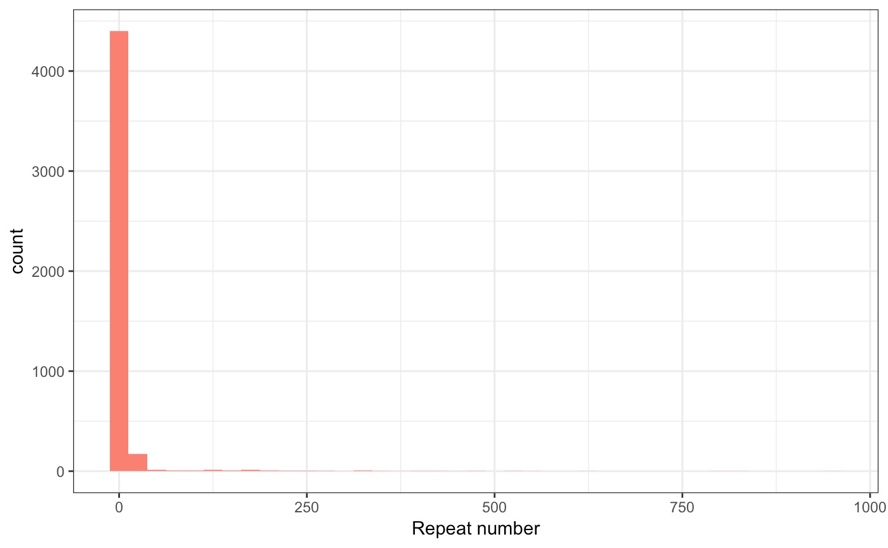

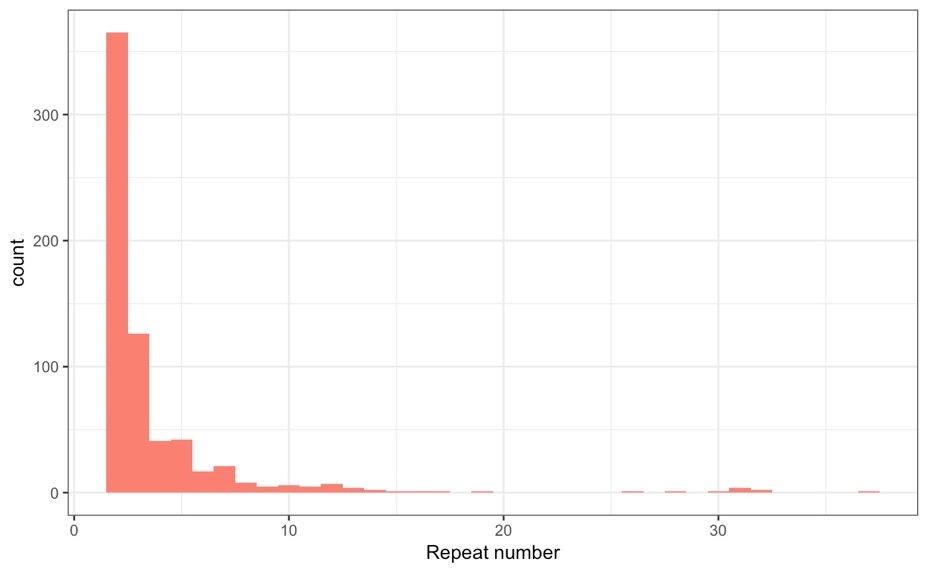

**Supplemental Figure 2. Most camouflaged regions are duplicated 2-3 times.** **(a)** We measured the number of times each gene region was duplicated and found that 70% of gene regions were replicated three or fewer times in the genome, but 84 regions were duplicated ≥100 times, with the most repeated region (intronic region from C5orf48) being replicated 941 times. **(b)** Limiting to only CDS regions, we estimate that 74.1% are replicated three or fewer times, with 38 replicated ≥10 times and the most repeated region was from NBPF12, which was replicated 37 times (Supplemental Figure 2b).

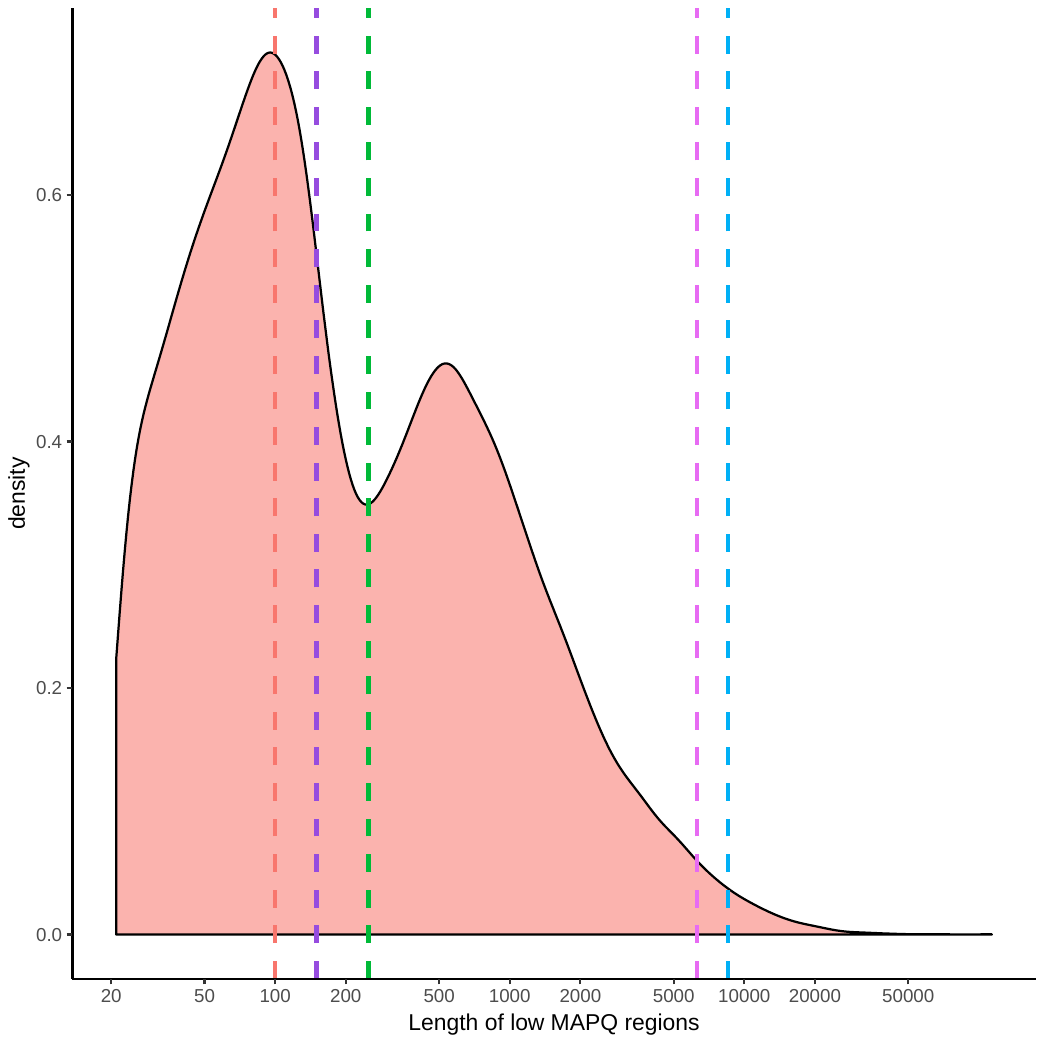

**Supplemental Figure 3. Distribution of camouflaged region sizes explains why increasing read lengths to 250 nucleotides and beyond resolves a large proportion camouflaged regions.** We generated a density plot for the length of all camouflaged regions to approximate the proportion of regions each sequencing technology should be able to resolve, which resulted in a bimodal distribution. The two modes are located at 95 and 538 nucleotides. As expected, median read lengths for the Illumina whole-genome sequencing based on 100-nucleotide (identified as the dashed red line) and 250-nucleotide (green dashed line) read lengths were 100 and 250 nucleotides, respectively. The first mode for the camouflaged region lengths is at 95, explaining why 100-nucleotide read lengths are insufficient to unambiguously span most dark-by-mapping quality regions. The 250-nucleotide read lengths fall between the two modes, explaining why 250-nucleotide read lengths resolve a high percentage of camouflaged regions. In other words, 100-nucleotide read lengths are too short to bridge most camouflaged regions, but 250-nucleotide read lengths appear to be sufficient for many. Median read lengths for both the ONT (magenta dashed line) and PacBio (blue dashed line) genomes we used in this study were 6276 (N50 = 33973) and 8511 (N50 = 17467) nucleotides, respectively, which is shorter than expected, but substantially longer than necessary to resolve most camouflaged regions. We believe comparing median read lengths, rather than N50, is more useful in this scenario, because we are interested to know what percentage of reads are likely to bridge a given dark or camouflaged region. Median 10x genomics read length (purple dashed line) is based on physical read length and not on synthetic read length.

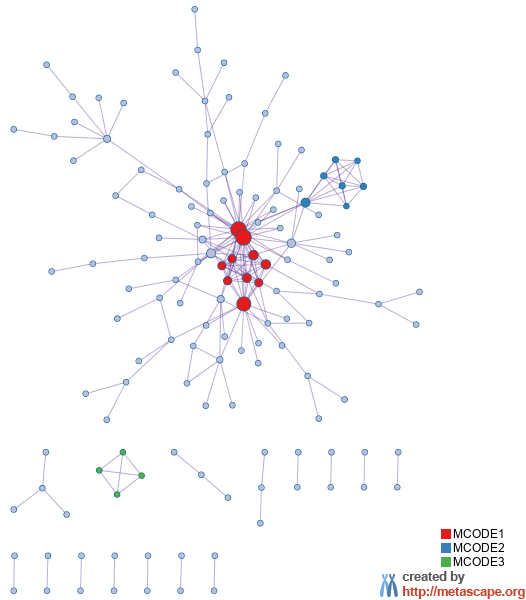

**Supplemental Figure 4. There are 212 known protein-protein interactions amongst 138 dark genes.** Looking specifically at known protein-protein interactions, we found 138 proteins with 212 known interactions.

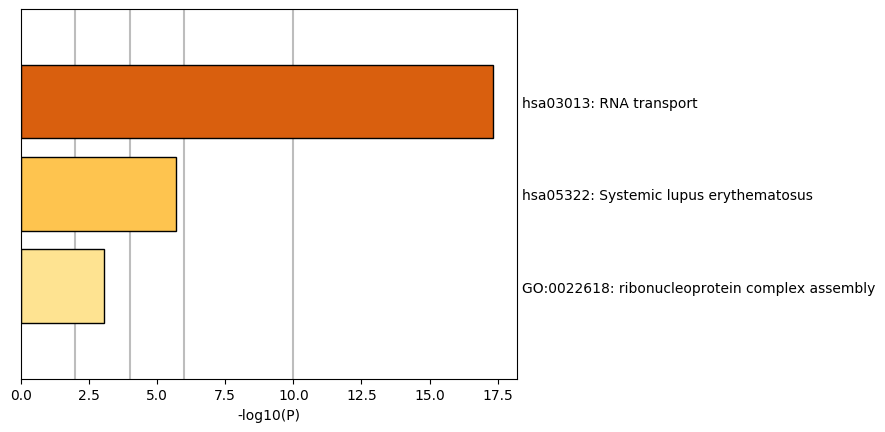

**Supplemental Figure 5. All three MCODE groups combined are primarily associated with RNA transport.**

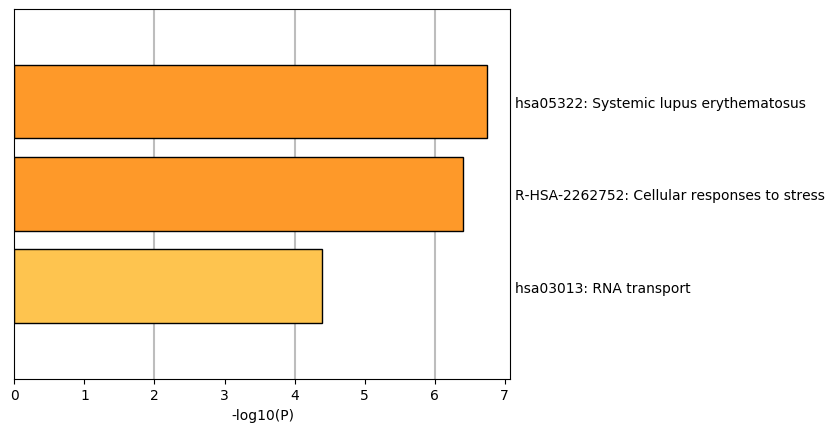

**Supplemental Figure 6. MCODE group 1 is enriched for systemic lupus erythematosus and cellular responses to stress.**

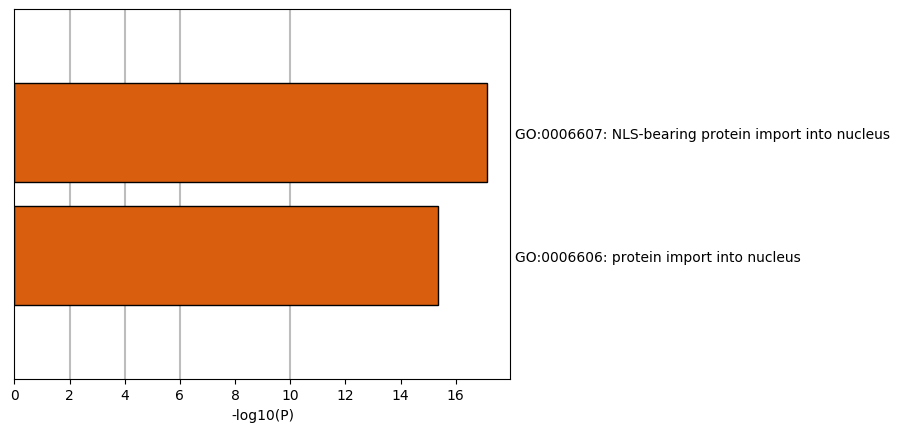

**Supplemental Figure 7. MCODE group 2 is enriched with proteins involved in NLS-bearing protein import into nucleus and protein import into nucleus**.

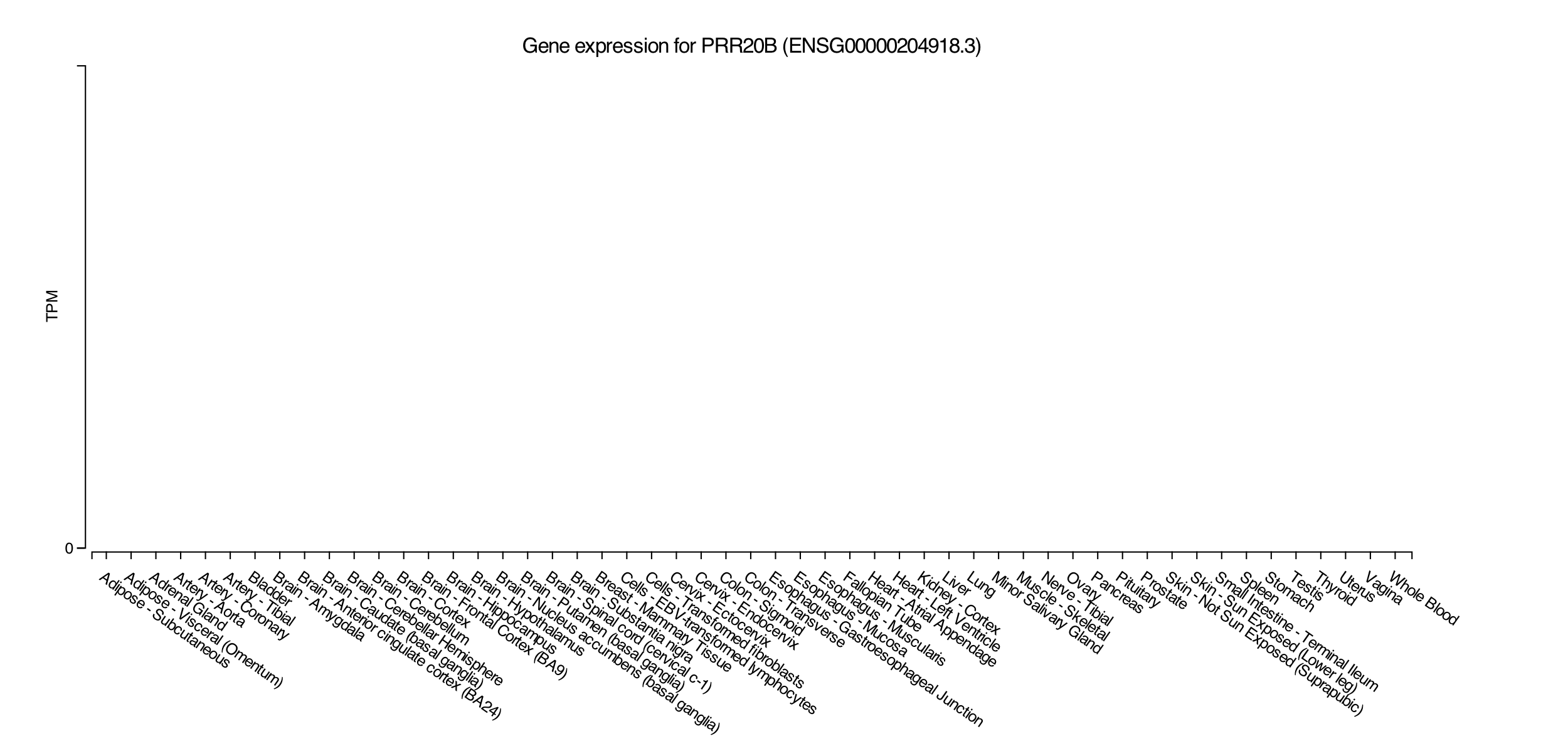

**Supplemental Figure 8. PRR20B is 100% camouflaged and has no known expression in GTEx (accessed December 2018).**

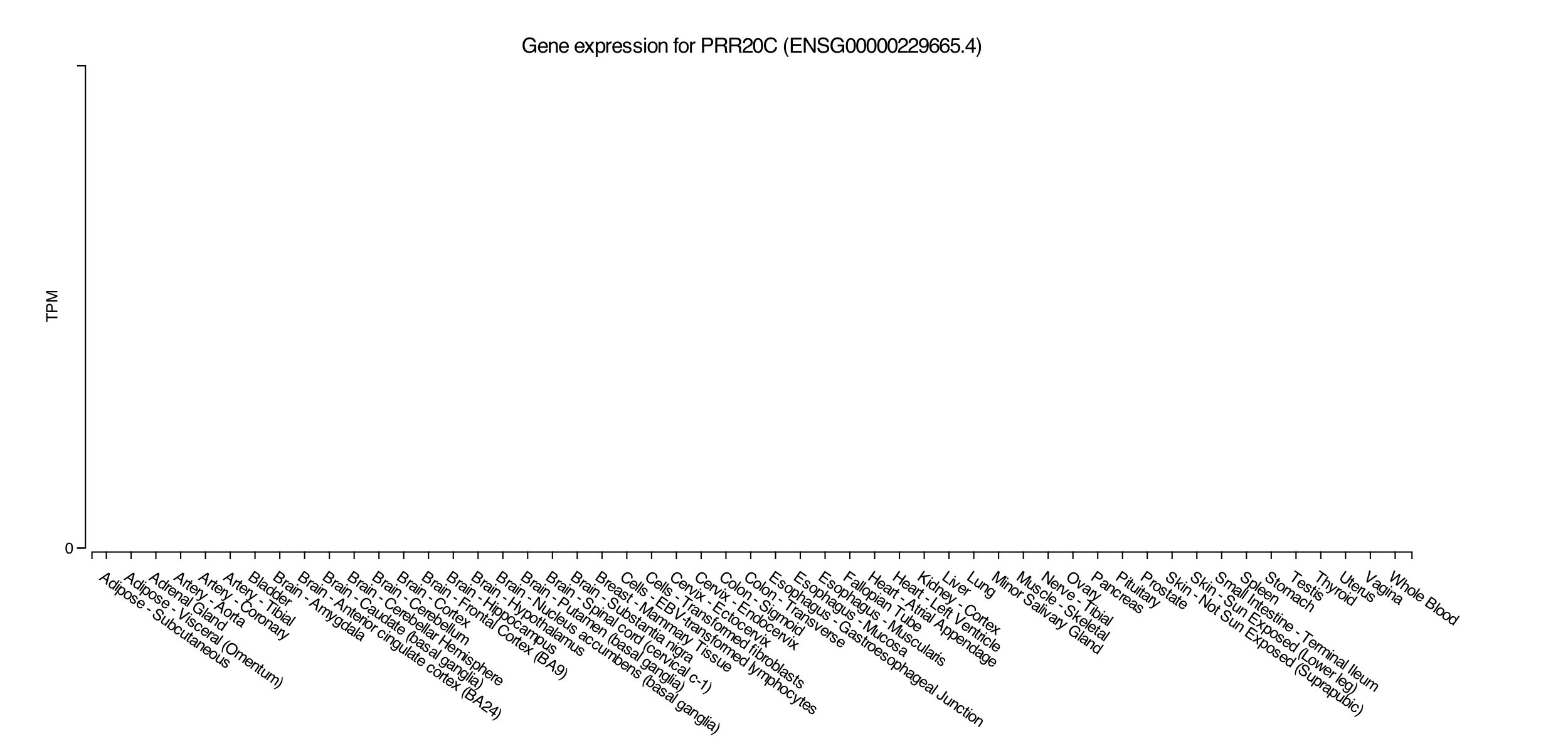

**Supplemental Figure 9. PRR20C is 100% camouflaged and has no known expression in GTEx (accessed December 2018).**

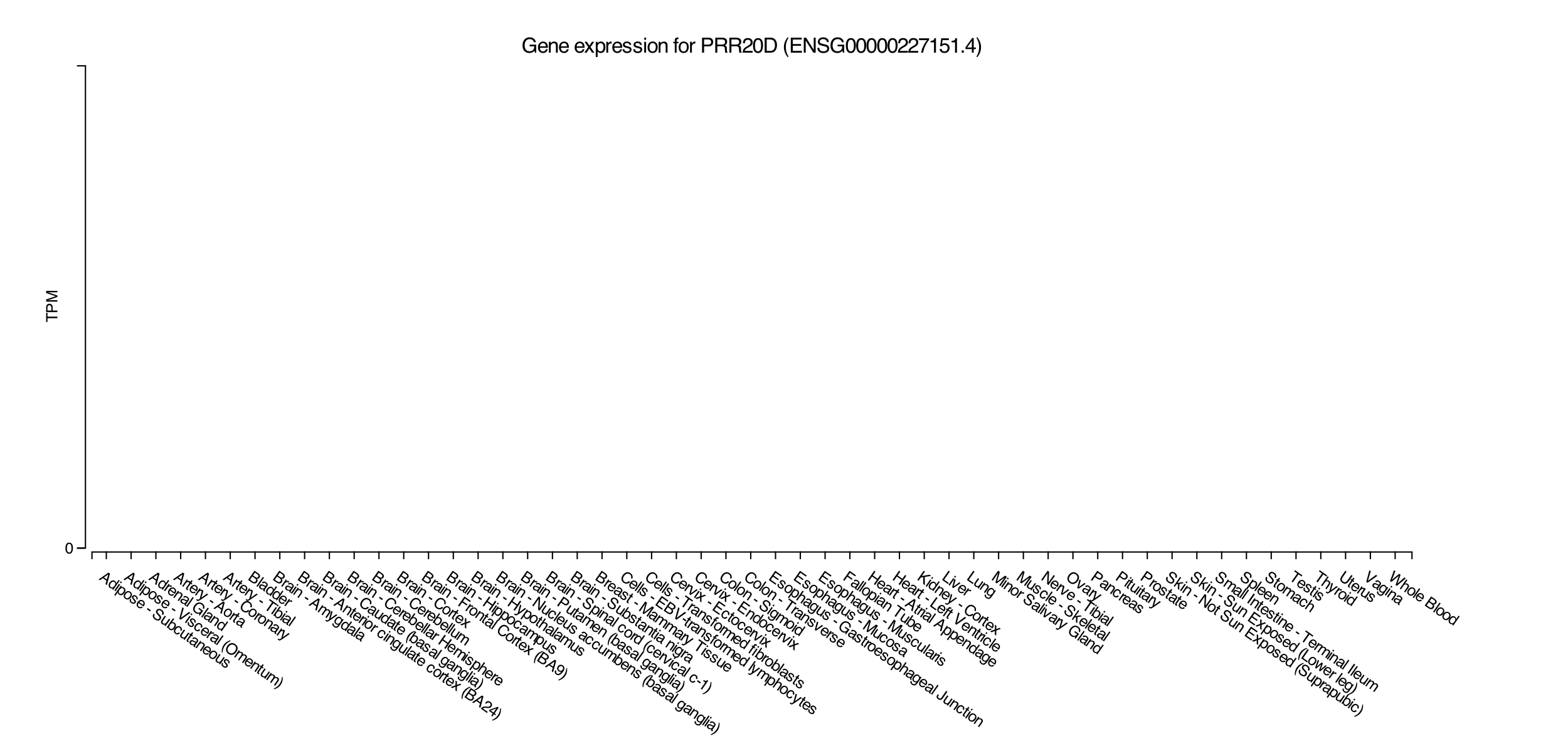

**Supplemental Figure 10. PRR20D is 100% camouflaged and has no known expression in GTEx (accessed December 2018).**

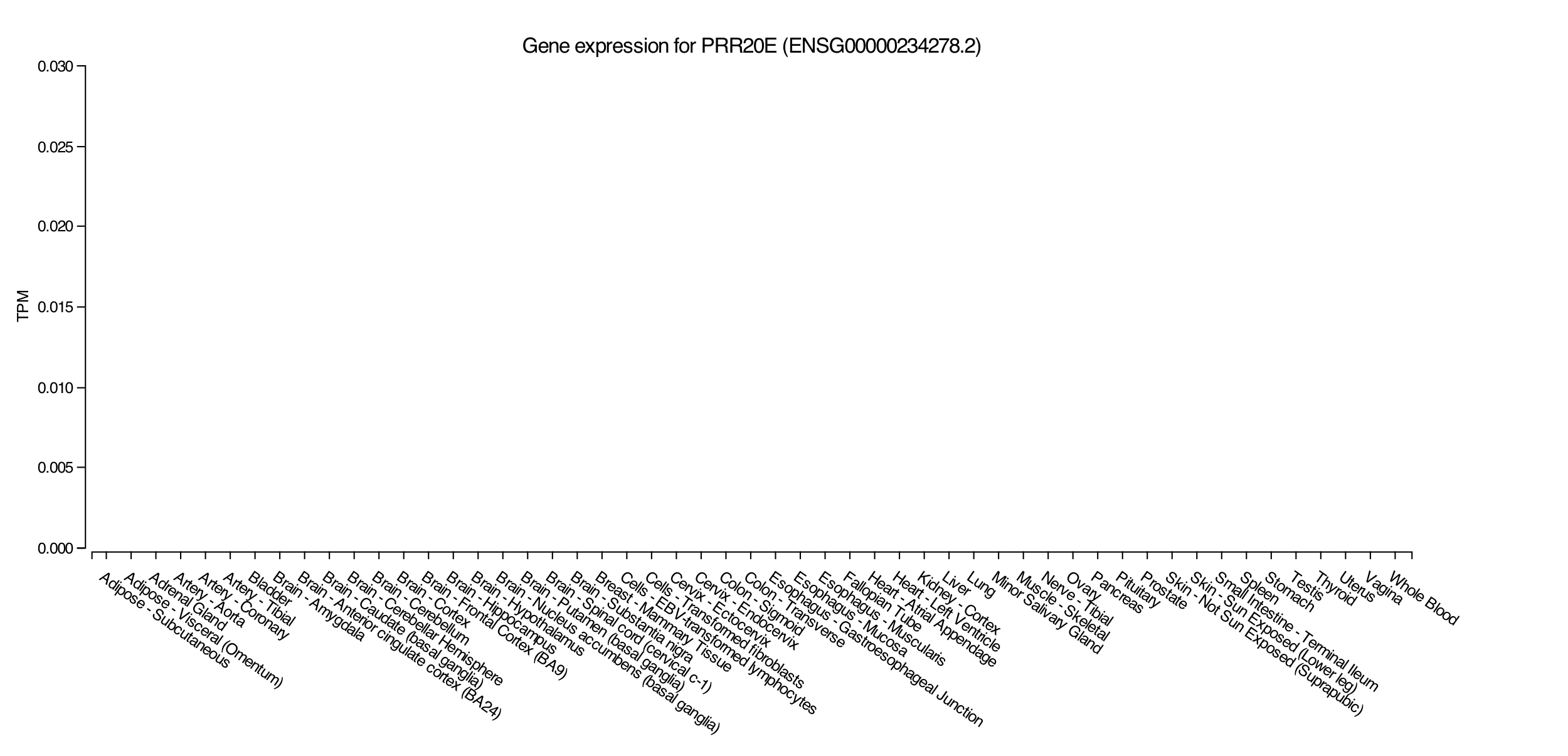

**Supplemental Figure 11. PRR20E is 100% camouflaged and has no known expression in GTEx (accessed December 2018).**

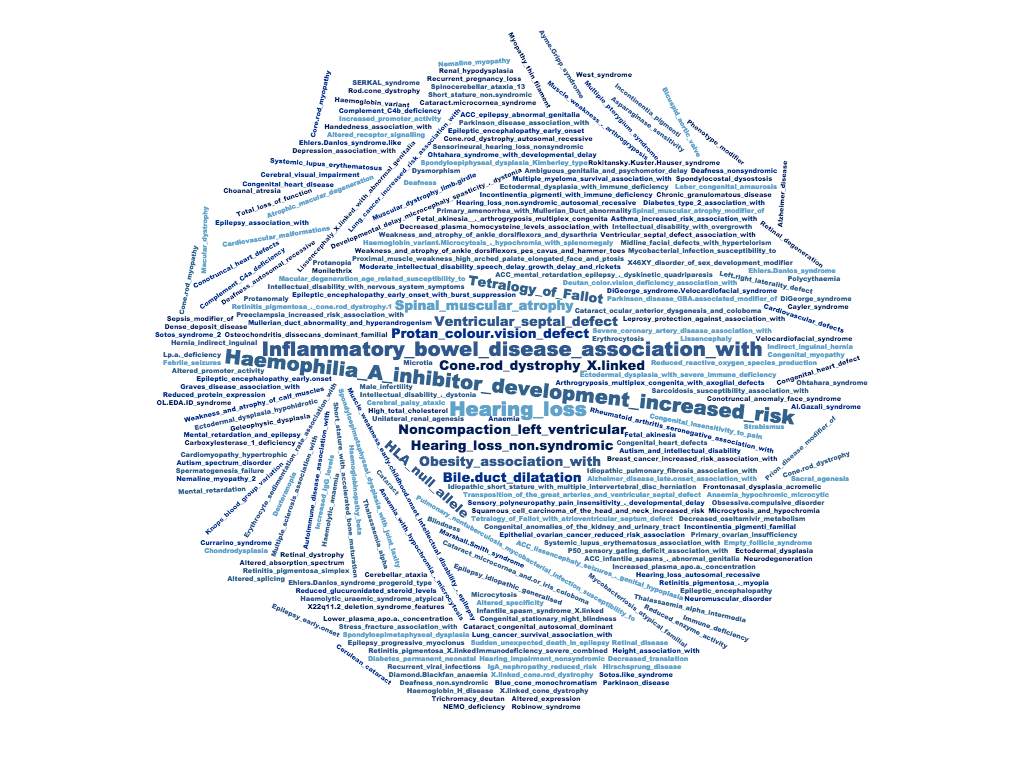

**Supplemental Figure 12. Dark genes are enriched for genes involved in several diseases, including Hemophilia A, color blindness (protan colour vision defect), and X-linked cone-rod dystrophy.** We performed an enrichment analysis, where the diseases most enriched for dark genes included Hemophilia A, color blindness (protan colour vision defect), and X-linked cone-rod dystrophy.

**APOE**

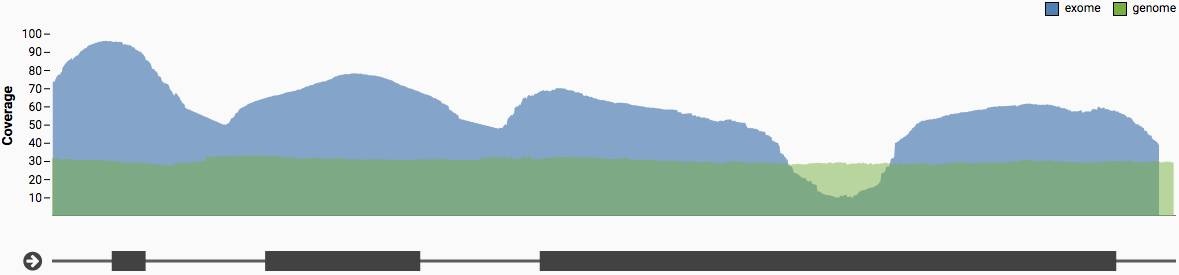

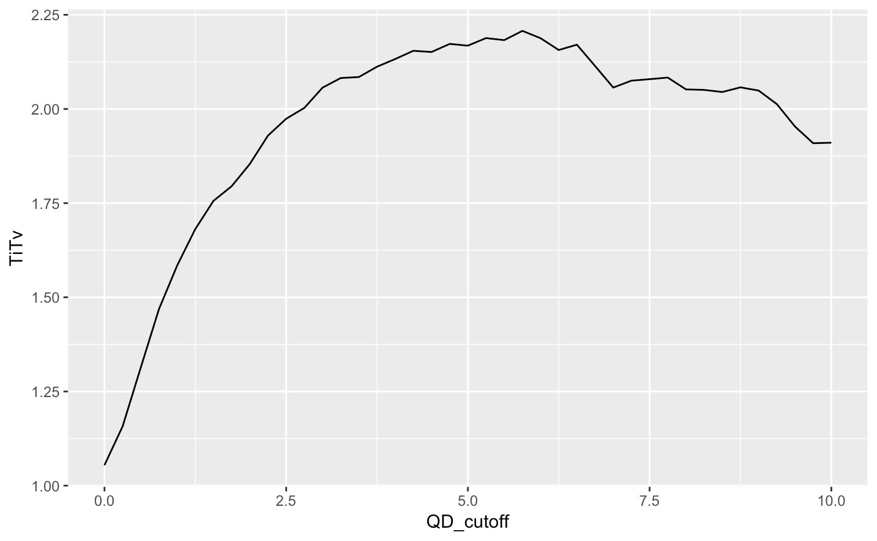
**Supplemental Figure 13. The *APOE* gene is 6% dark in whole-exome data, and is dark for some samples with whole-genome sequencing.** *APOE*—the top genetic risk for Alzheimer’s disease—is approximately 6% dark CDS (by depth) for certain ADSP samples with whole-genome sequencing, and the same region is dark in gnomAD whole-exome data.

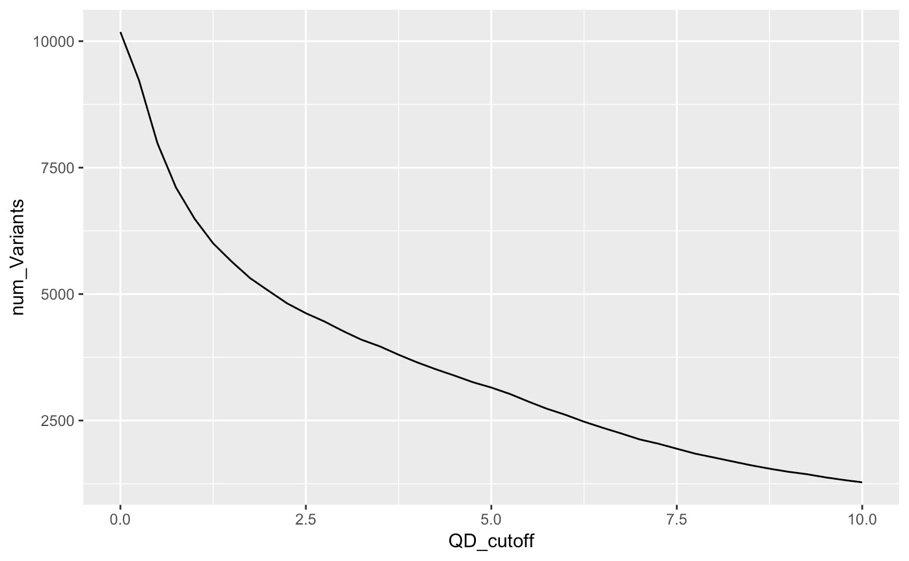

**Supplemental Figure 14. Our method rescues between approximately 3152 and 4622 variants in 13142 ADSP samples.** Across 13142 samples from the ADSP, excluding all variants with a quality by depth (QD) <2.5, were able to rescue approximately 4622 exonic variants with a transition-transversion ration (Ti/Tv) of 1.97 from 147 camouflaged region sets, that are spread across 501 camouflaged genes (Supplemental Figure 14; Supplemental File 1). Using more stringent QD (excluding variants with QD <5), we rescued 3152 variants with a Ti/Tv ratio of 2.17. We only included exons from genes that are at least 5% dark CDS. **(a)** We plotted Ti/Tv ratios versus QD cutoff. The Ti/Tv ratio at QD = 2.5 and QD = 5.0 are 1.97 and 2.17, respectively. **(b)** The number of variants rescued at QD = 2.5 and QD = 5 are 4622 and 3152, respectively.
